# The ancient evolution of far-red light photoacclimation in cyanobacteria

**DOI:** 10.1101/2024.05.15.594131

**Authors:** Laura A. Antonaru, Cecilia Rad-Menéndez, Susan Mbedi, Sarah Sparmann, Matthew Pope, Thomas Oliver, Shujie Wu, David Green, Muriel Gugger, Dennis J. Nürnberg

**Affiliations:** Department of Life Sciences, Imperial College London, London, UK; Institute for Experimental Physics, Freie Universität Berlin, Berlin, Germany; Culture Collection of Algae and Protozoa, Scottish Association for Marine Science, Oban, UK; Berlin Center for Genomics in Biodiversity Research, Berlin, Germany; Museum für Naturkunde, Leibniz Institute for Evolution and Biodiversity Science, Berlin, Germany; Leibniz Institute for Freshwater Research and Inland Fisheries, Berlin, Germany; Esox Biologics, London, UK; Department of Physics and Astronomy, Vrije Universiteit Amsterdam, Amsterdam, the Netherlands; Dahlem Centre of Plant Sciences, Freie Universität Berlin, Berlin, Germany; Institut Pasteur, Université Paris Cité, Collection of Cyanobacteria, Paris, France

**Keywords:** Cyanobacteria, oxygenic photosynthesis, photoacclimation, FaRLiP, chlorophyll f, microbialites, microbial mat, phylogeny, evolution

## Abstract

Cyanobacteria oxygenated the atmosphere of early Earth and continue to be key players in global carbon and nitrogen cycles. A phylogenetically diverse subset of extant cyanobacteria can perform photosynthesis with far-red light through a process called far-red light photoacclimation, or FaRLiP. This phenotype is enabled by a cluster of ∼20 genes, and involves the synthesis of red-shifted chlorophylls *f* and *d*, together with paralogues of the ubiquitous photosynthetic machinery used in visible light. The FaRLiP gene cluster is present in diverse, environmentally important cyanobacterial groups but its origin, evolutionary history, and connection to early biotic environments have remained unclear. This study takes advantage of the recent increase in (meta)genomic data to clarify this issue; sequence data mining, metagenomic assembly, and phylogenetic tree networks were used to recover more than 600 new FaRLiP gene sequences, corresponding to 52 new gene clusters. These data enable high-resolution phylogenetics and - by relying on multiple gene trees, together with gene arrangement conservation - support FaRLiP appearing early in cyanobacterial evolution. Sampling information shows that considerable FaRLiP diversity can be observed in microbialites to the present day, and the process may have been associated with microbial mats and stromatolite formation in the early Paleoproterozoic. The ancestral FaRLiP cluster was reconstructed, revealing a conserved intergenic regulatory sequence that has been maintained for billions of years. Taken together, our results indicate that oxygenic photosynthesis using far-red light may have played a significant role in Earth’s early history.

## Introduction

Cyanobacteria are important primary producers, and they have played a fundamental part in the Great Oxygenation Event that changed Earth’s atmosphere^1–3^. The evidence can be seen in fossil stromatolites, some of the earliest evidence we have of life on Earth^2,4^. Molecular clock data combined with the fossil record approximates their last common ancestor at 2.3 - 3.4 billion years ago^5–8^. Since then, cyanobacteria have diversified, colonizing a wide variety of habitats from the low-nutrient open ocean to the interior of rocks in hyperarid deserts, including regions inhospitable to eukaryotic photosynthetic organisms^3,9^. Some of this environmental diversity is reflected in morphology, as cyanobacteria encompass lineages that are unicellular, or filamentous, including those with specialized nitrogen-fixing cells^7,10^. Filamentous growth forms have evolved multiple times, and this morphology may have been acquired early on in cyanobacterial evolution^7,11^.

Filamentous cyanobacteria are vital for understanding one of the major early biological innovations on Earth: the development of microbial mats. These structures would have protected against UV-C radiation and, through calcification and sedimentation, further led to the formation of stromatolites. These were major biotic environments for a period of ca. 2 billion years during the Proterozoic^7,12^. Based on data from extant early lineages, the earliest cyanobacterial morphotypes likely had small cell diameters (<2.5 µm)^5,6^. Despite their evolutionary importance, few strains and genomes are available for early, thin-filamentous cyanobacteria^13^. This leaves gaps in the data when investigating, for example, the evolution of ancient shading responses concomitant with the emergence of stromatolites.

Photosynthetic organisms can adapt to different light conditions. Diverse mechanisms have evolved for photoacclimation since the start of photosynthesis, and organisms may change their photosynthetic systems in response to light quantity or quality (e.g. spectral niches)^14–17^. Spectrum-based photoacclimation can help cyanobacteria to thrive in various niches, from clear open ocean (enriched in blue light) to increasingly turbid seas and coastal waters (red light), as well as microbial mats and terrestrial environments (far-red light)^15,18^. Patterns have been observed on a global scale, and some types of photoacclimation are present in many ancient lineages^19–21^.

One recently-discovered spectral adaptation is Far-Red Light Photoacclimation (FaRLiP). Its evolution is still unclear. This process relies on red-shifted chlorophylls, chlorophyll *d* and chlorophyll *f*, in order to allow photosynthesis in the near-infrared / far-red (700-750 nm)^22–25^. The distribution of FaRLiP cyanobacteria appears to favor shaded environments, which are enriched in far-red light^16^. These include microbial mats, the subsurface of rocks, and microbialites (sedimentary structures formed by microbes), such as present-day stromatolites and thrombolites^16,23,26–32^.

The basis of this adaptation is a cluster of around 20 genes, encoding not only the enzyme synthesizing chlorophyll *f* (*chlF*), but also alternative components of Photosystem I (*psa)*, oxygen-evolving Photosystem II (*psb*), phycobilisomes (*apc*), and a phytochrome signaling cascade (*rfp*)^22,33–39^. Across the cyanobacterial species tree, FaRLiP is rare, occurring in a small subset of morphologically and phylogenetically diverse cyanobacteria, including early-branching filamentous lineages. For this reason, it was originally hypothesized to have been largely transmitted by horizontal gene transfer (HGT)^37^. However, recent work on ApcE2, the far-red phycobilisome-membrane linker, has emphasized the significance of vertical descent combined with gene loss in driving the evolution of this adaptation^26^. This hypothesis has been generally accepted^39–42^. Nevertheless, the data was insufficient to fully support vertical descent from the cluster’s Most Recent Common Ancestor (MRCA).

Early horizontal gene transfer between distantly related cyanobacteria remained a possibility. Trees of different FaRLiP genes were not fully congruent^37,38^, and considerable variation existed in gene arrangement^24,43^, raising the possibility of extensive single-gene HGT. In particular, the HGT of Photosystem I subunits to early FaRLiP lineages has been proposed^38^. The limited FaRLiP data from early filamentous lineages was a particularly large issue^26^. It left the early evolutionary history of FaRLiP unclear. However, FaRLiP gene fragments associated with early lineages had been recovered from the Sequence Read Archive (SRA)^26^.

This work aims to clarify the evolution of FaRLiP. Sequence data mining of genomes and metagenomes ensured the recovery of many novel, highly diverse FaRLiP gene clusters. A comprehensive phylogenetic analysis was performed, combining gene trees, genome trees, synteny, GC content analysis and metatranscriptomics, as well as investigations into intergenic regions. We demonstrate that there is strong evidence for the cluster being vertically inherited since its MRCA, with losses in many cyanobacterial groups, and the maintenance of a conserved regulatory motif since its origin.

## Results

### Far-Red Light Photoacclimation is associated with microbialites and microbial mats

A total of 52 new complete or nearly complete FaRLiP gene clusters were recovered from a variety of sources. These include the NCBI database (14 clusters), additional cultured strains (16), raw SRA sequence data that was assembled in the course of this project (20), and enriched isolates from our laboratory (2) (Tables S1-S4). The significance of metagenomic data as a resource for this project cannot be overstated. More than half of the clusters were sourced from metagenome-assembled genomes (MAGs) from species-rich environmental samples. Overall, common sampling locations included microbialites - both stromatolites and thrombolites (24%). FaRLiP was also present in non-lithifying microbial mats (23%), together with other types of environments (48%) and unknown sources (5%) (Figure 1A).

**Figure 1.**
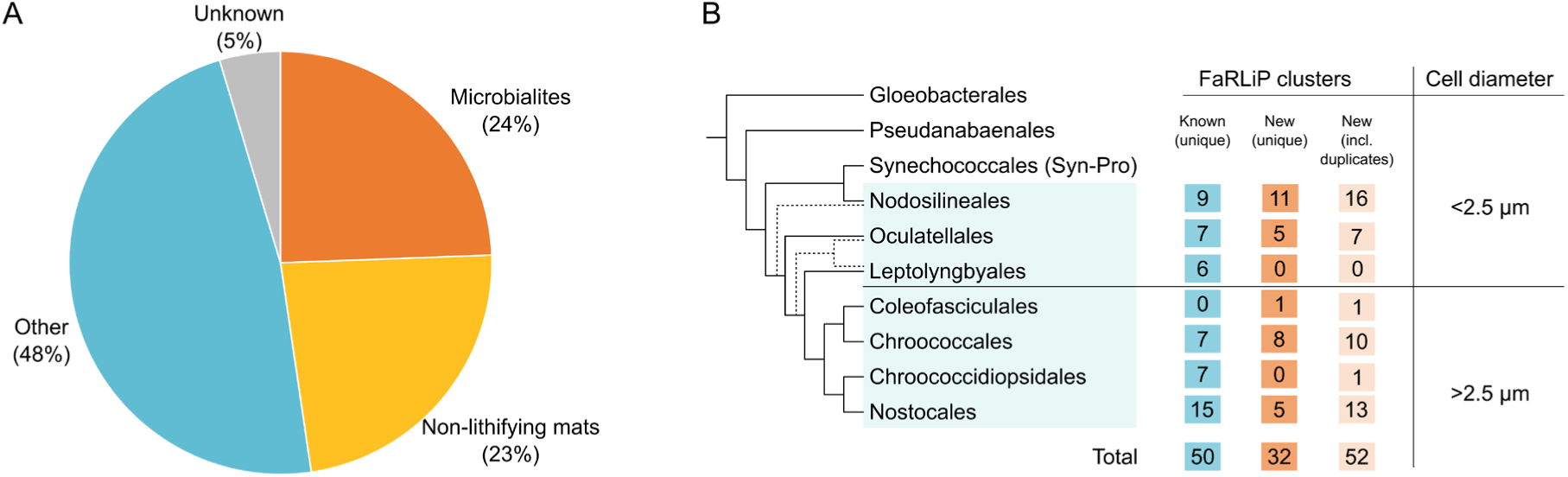
The environmental and phylogenetic distribution of FaRLiP. (A) Microbialites, together with non-lithifying mats, are the source of about half of the FaRLiP gene clusters identified in this work. Environmental dataset size: 86 clusters. (B) FaRLiP is present in a wide range of cyanobacterial orders indicated by a blue-green box. The relationships between these lineages are shown on the left; dashed lines mark alternative phylogenetic paths. The table shows the number of clusters associated with each order, already known (in blue) or newly identified, counted either as unique sequences (orange) or including duplicates (light orange).

The microbialite sampling sites varied geographically and environmentally, encompassing five coastal and lake locations from three continents. Two of the stromatolite sites, notably Alchichica Lake (Mexico) and Cape Recife/Schoenmakerskop (South Africa) showed remarkable FaRLiP diversity. They contained ≥6 FaRLiP clusters (and MAGs) each, from four cyanobacterial orders, and approximately half of the cyanobacterial MAGs in these datasets had FaRLiP genes. Given the rarity of microbialites in the current Phanerozoic eon, the presence (and occasional abundance) of FaRLiP genes in so many of them suggests a significant role for FaRLiP in this habitat.

The only other microenvironments with a similarly high diversity and abundance of FaRLiP genes were marine microbial mats on microplastics in the Pacific Garbage Patch. FaRLiP gene fragments have been reported from there before, and were presumed to be associated with mat-forming, filamentous cyanobacteria inhabiting the plastics^26,44,45^. Microplastics-associated sequences were vital for recovering data from previously underrepresented, early-branching lineages of thin (cell diameter <2.5 µm), filamentous cyanobacteria^6^, such as *Nodosilineales* and *Oculatellales* (Figure 1B), and helped to resolve early FaRLiP evolution.

### FaRLiP motifs improved data recovery

Relying on metagenomic data has its challenges. Rarely, the resulting clusters were present within large (>100 kbps) contiguous sequences within the metagenomes, but often, the FaRLiP gene clusters were fragmented. Therefore, it was not sufficient to use just a single motif from the ApcE2 sequence as in previous work^26^. Far-red motifs for additional genes were developed as search queries in order to recover more sequences for later phylogenetic work. Previous publications have looked at amino acids unique to FaRLiP paralogues^21,36–39^. Occasionally these could be used directly (such as the QD site in ChlF)^36^, but sometimes these residues proved impractical as queries due to lack of conservation in neighboring amino acids (e.g. the FaRLiP-specific PsaA2 loop)^38^. In this study, the analysis was done by studying amino acid alignments for areas that were conserved in FaRLiP paralogues, but distinct from white-light versions (Figure 2). Many of these conserved regions have known or hypothesized functions related to far-red light photoacclimation (Table S5). Overall, >600 new FaRLiP genes were recovered, enabling for the first time more complex analyses of ancient FaRLiP evolution.

**Figure 2.**
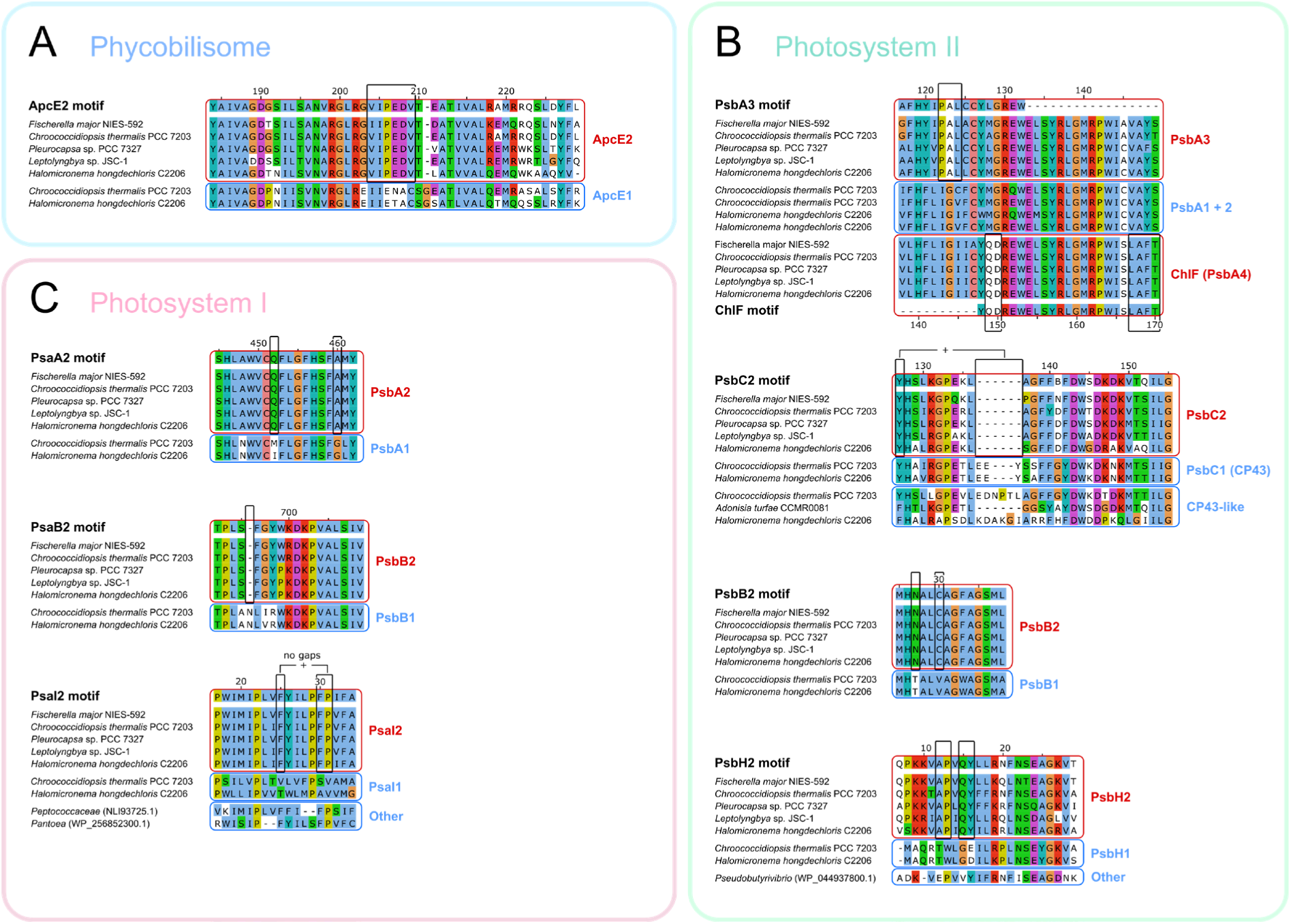
FaRLiP paralogues show conserved amino acid motifs. They can be observed in phycobilisome (A), Photosystem II (B) and Photosystem I (C) proteins. These motifs are conserved between FaRLiP sequences in distantly related species (red boxes), and also easily distinguished from standard/white light paralogues or other similar sequences (blue boxes). For PsbA, there are two FaRLiP paralogues: PsbA3 (far-red D1), and ChlF (also known as PsbA4; the chlorophyll *f* synthase). Black vertical boxes mark the main distinguishing features. For most genes, any single feature shown was sufficient to identify FaRLiP paralogues. For PsaI2 and PsbC2, multiple features had to be considered together. See also Table S5.

### The evolution of FaRLiP genes supports vertical descent

The FaRLiP cluster consists of 19-24 genes. Out of them, 19 core genes are present in all known cyanobacteria with a complete FaRLiP cluster. They encode paralogous components of phycobilisomes (*apcE2/D2/D3/D5*), Photosystem I (*psaA2/B2/I2/L2/F2/J2*), Photosystem II (*psbA3/B2/C2/D3* and *chlF*), and a phytochrome signaling cascade (*rfpA/B/C*). Their functions have been described in detail before^22,24,33,35,36^. Phylogenetic trees were inferred from all the 19 core genes (Table S6). The tree for *apcE2*, the gene encoding the phycobilisome-photosystem linker, was especially highly supported. It also accurately matched the species tree (Figure 3). This provides strong evidence for vertical descent from a common ancestor, and, due to the rarity of FaRLiP, implies that there were repeated losses of the gene cluster throughout evolutionary history. Sequences for *apcE2* fall into six groups, equivalent to seven orders: *Nostocales* (I), *Chroococcidiopsidales* (II), *Coleofasciculales* (III), *Chroococcales* (IV), *Oculatellales* and *Leptolyngbyales* (V) and *Nodosilineales* (VI). Group V includes two orders that could not be reliably separated, as the branching of the FaRLiP sequences within them was not well resolved. Out of all groups, *Coleofasciculales* (III) is new to FaRLiP research. It is represented by a single strain, *Wilmottia* sp. PCC 9708.

**Figure 3.**
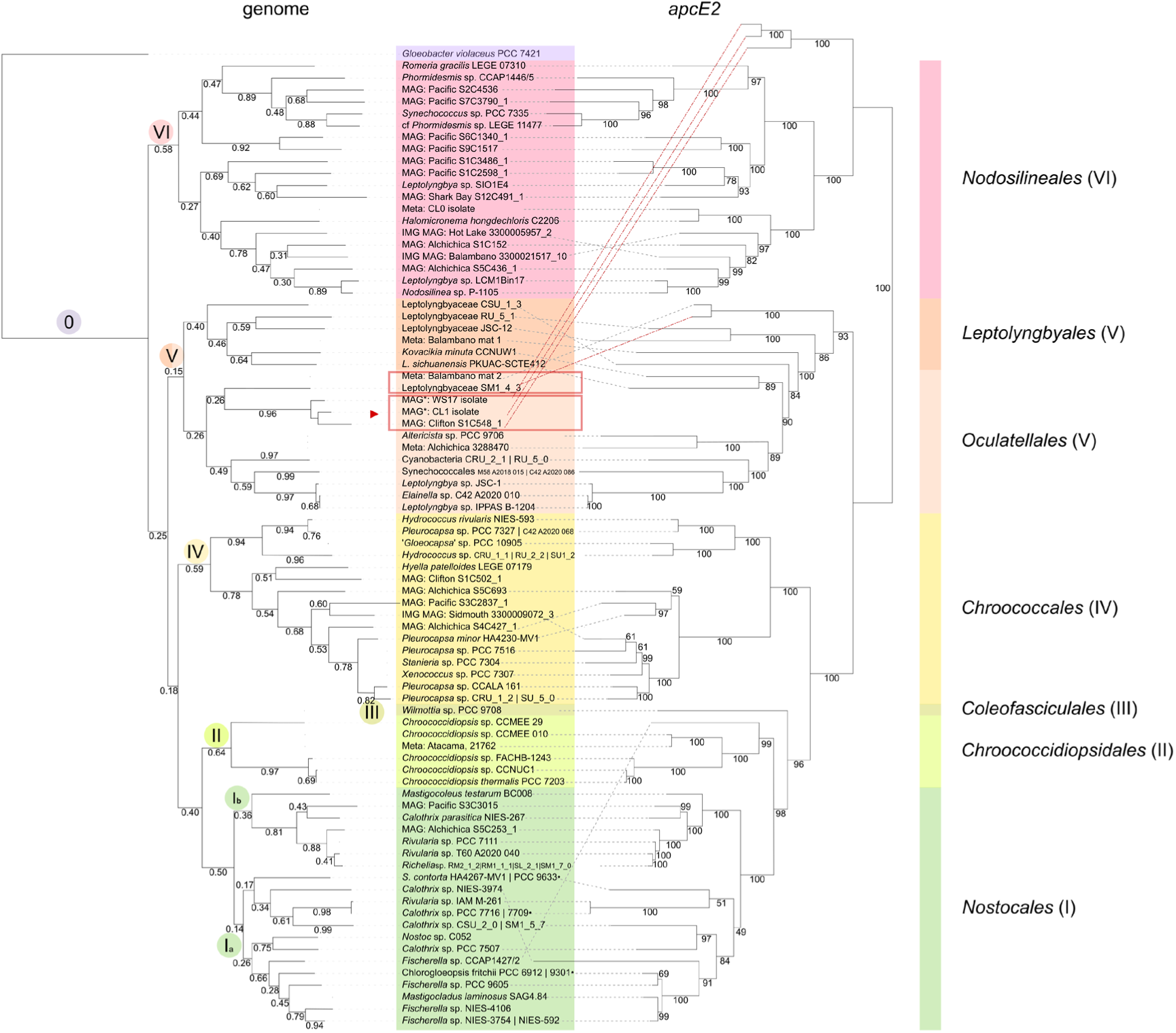
The genome tree (left) resembles the tree of a FaRLiP marker gene (*apcE2*, right). This suggests that the FaRLiP cluster is an ancient innovation that mainly evolved by vertical descent. It is present in seven orders, listed on the right, organized into six groups (I-VI). On the left, in circles, the genome tree is cross-referenced with the different cluster orientations in Figure 4. Red boxes in the middle mark branches where FaRLiP gene phylogenies are different from the genome tree. *Gloeobacter* (purple) was used as outgroup. Bootstrap 100.

All core FaRLiP gene trees were generally congruent with each other, as well as the species tree (Table S6). The same six groups (I-VI) were often present. However, some discrepancies remain. In previous work it was uncertain whether they were caused by biological factors (e.g. HGT) or by technical artefacts (e.g. different evolutionary rates, limited data)^37,38^. The present study lends strong support for the latter. More similar gene trees were observed in this study overall, likely due to the increased data availability.

The dissimilarities of greatest interest were those in which the gene trees agreed between themselves, but not with the species tree, as this may indicate HGT of the whole gene cluster. Two branches of the *Oculatellales* (V) show such incongruities (red boxes in Figure 3). In one case, FaRLiP sequences of MAG origin were assigned to a different order (VI) than the genome (V) (red arrow in Figure 3). This is unlikely to be due to MAG contamination, as the three genomes involved come from two separate geographic locations and sequencing projects. While HGT cannot be fully excluded, it is more likely that this incongruity represents a Long Branch Attraction artifact instead. Between the aforementioned entries and known homologues, there is low sequence similarity (AAI: 73%) at both the genome and the cluster level. Large evolutionary distances make phylogenetic reconstruction more prone to artifacts^46^, and as such, vertical descent remains the most likely possibility.

The wide range of evolutionary rates and sizes between FaRLiP genes also explains most of the differences remaining between the gene trees (Figure S1, Table S7). Large genes (> 300 bp), that evolve moderately fast, such as phycobilisome and Photosystem II (PSII) genes, tended to also have gene trees that agreed in fine detail, and agreed with the species tree. This can be visualized by ‘overlapping’ the trees into networks (Figure S2). To a lesser extent, this agreement was also observed for the faster-evolving *rfp* genes of the phytochrome signaling cascade (Figure S2D). In contrast, most mismatches were observed in Photosystem I (PSI) gene trees.

It has been previously suggested that these mismatches indicate different phylogenetic histories, with PSI genes evolving later and then being transferred by HGT to early branches of the FaRLiP cyanobacterial tree^38^. This hypothesis discounts the influence of divergent evolutionary rates on phylogenetic reconstruction. In particular, FaRLiP PSI genes include both very small, fast-evolving genes such as *psaI*/*J*, and the very slow-evolving *psaA*/*B* (Table S7). We demonstrate this effect by performing a minor artificial alteration of reverse-complementing sequences prior to alignment and tree-building (the ‘Heads or Tails’ method, or HoT)^47^. This significantly changed the shape of the aforementioned PSI gene trees, due to unreliable alignments, but not of the others. Therefore, when evolutionary rates are taken into consideration, all core FaRLiP genes appear likely to share a common evolutionary trajectory, suggesting they have been inherited together.

Apart from the 19 core genes that are present in all clusters, other FaRLiP paralogues may occur in specific lineages. Previous research has identified three of them through experimental data (*psbH2*/*O2*/*V2*)^22,48–51^ and another through bioinformatics^39^ (named here *psbH2’*). This study made it possible to recover two additional genes, *psbF2* (a minor PSII gene) and *rfpD* (a putative regulatory protein) (Table S8). Although the small size of these genes severely limits the accuracy of phylogenetic trees, the overall pattern mirrors the one of the core genes.

Overall, the evidence supports the co-inheritance of all genes in the FaRLiP cluster from a common ancestor. This MRCA would have existed prior to the split of the earliest FaRLiP-containing cyanobacterial lineage, *Nodosilineales* (VI), from the cyanobacterial species tree. Early cyanobacterial lineages in which FaRLiP is present (V-VI) have previously been associated with evolutionary innovations related to microbial mat formation, such as nitrogen fixation, extracellular polysaccharide sheath formation and, in some cases, larger cell sizes^5–7^.

### FaRLiP cluster architecture reveals evolutionary patterns

FaRLiP gene arrangement (synteny) is conserved in a manner consistent with the genome tree, which further supports vertical descent of this phenotype. Additional evidence is provided by GC% values that are very similar between gene clusters and associated genomes (Table S9). Synteny was used to classify FaRLiP clusters, which also resolved the same six groups (I-VI) as sequence similarity did. There is typically one major (most frequently encountered) syntenic variant of the FaRLiP cluster in each group (Figure 4A). All these variants all share certain conserved characteristics in gene arrangement, but also exhibit aspects that are group specific.

**Figure 4.**
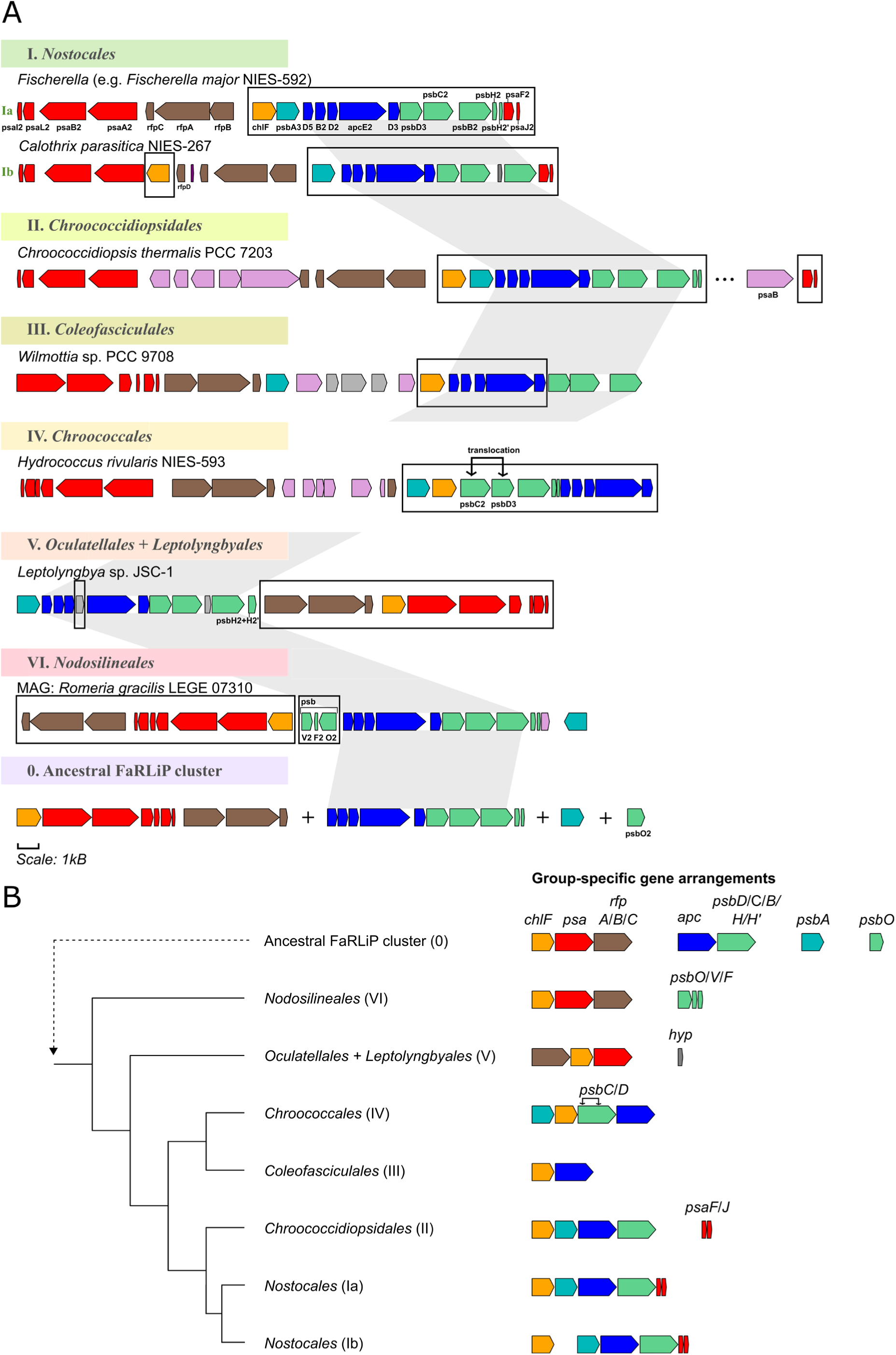
Synteny and gene orientation is conserved within FaRLiP gene clusters, in a manner consistent with vertical descent. (A) There are six cyanobacterial groups in which FaRLiP is present (I-VI). Each shows gene arrangements that are specific to itself (black boxes) as well as one feature that is conserved between most orders (*apc*, followed by *psb* genes, grey highlight). Only the most abundant syntenic variants are listed here. Based on both abundant and rare variants (Figure S3), the composition of the ancestral FaRLiP cluster (0) may be inferred. (B) Syntenic features specific to each group (right) are conserved in a way consistent with vertical descent (species tree schematic shown to the left). Color code: PSI (red), PSII (green), apart from *psbA3* (turquoise), *chlF* (yellow), phycobilisome (blue), regulatory signaling cascade (brown), non-FaRLiP hypothetical proteins (gray), and other proteins (pink).

Gene arrangements that are (near-)universal were likely present in the ancestral FaRLiP cluster. This includes the order of *apc* genes, followed by *psbD/C/B*, which is common across all lineages (except IV) (Figure 4A). In some cases, *psbA3* may be present upstream (I, II, V). This pattern suggests that co-transcription and associated co-regulation, especially between phycobilisome and PSII genes, are important for FaRLiP. Linkage can also be seen between the PSI genes (*psaA*/*B*/*L*/*I*/*F*/*J*). These are always found together in the same orientation (apart from minor components such as *psaF* and *psaJ*). Phytochrome signaling cascade genes (*rfpA*/*B*/*C*) are also always syntenically linked, with the exception of the optional downstream component *rfpD*.

Other gene arrangements are group-specific, and conserved in a manner consistent with vertical descent. This includes: *psaF/J* being present at the opposite end of the cluster from other PSI genes (in group I) or even in another area of the genome (group II); *chlF* immediately upstream of *apc* genes (III); a unique translocation of *psbC*/*psbD* (IV); the order of *rfp*, *chlF*, then *psa* (V) or, in contrast, *chlF*, *psa*, then *rfp* (VI) (Figure 4A, B). Some groups (I, II) exhibit only minor variants, both within and between them (Figure S3, S4). Others, especially early lineages (V, VI) show a remarkable diversity, with evidence for multiple events of cluster splitting, gene translocation and inversion (clusters 26, 29, and 30 in Figure S3).

Groups V and VI represent the earliest FaRLiP lineages to branch out from the cyanobacterial species tree (Figure 4B). Aspects that they share in common may define ancestral FaRLiP traits. The genes *rfp*, *psa*, and *chlF* are typically grouped together, though in different orientations. Which orientation is the oldest might be revealed by a minor cluster variant in lineage V (clusters 24-25 in Figure S3; red box marked with an arrow in Figure 3). It shares with other *Oculatellales* (V) the presence of a conserved hypothetical protein (Figure 4A). However, just like the earlier-branching *Nodosilineales* (VI), it contains the gene orientation of *chlF*, *psa*, then *rfp*, as well as the optional FaRLiP gene *psbO2*. The standard, white-light version of this gene is known to stabilize the manganese cluster of Photosystem II. The presence of *psbO2* may represent an ancestral trait.

The cluster in the most recent common ancestor of known FaRLiP cyanobacteria possibly had the following four elements. 1) It co-transcribed *chlF* with downstream PSI genes. Regulatory *rfp* genes would be further downstream. 2) Allophycocyanin genes would be co-transcribed with downstream PSII genes, which included minor genes *psbH2* and *psbH2’*. 3) The FaRLiP D1 gene, *psbA3*, would have its own promoter. 4) Potentially, this organism would have a far-red *psbO* and, tentatively, a far-red *psbV* and *psbF*, as seen in the earliest lineage, VI (Figure 4).

### The FaRLiP genes are co-regulated in operons through an ancient motif

An important aspect of the evolution of FaRLiP is its regulation. The highly conserved nature of FaRLiP gene arrangements suggests that parts of the cluster could be co-transcribed as operons. This hypothesis is supported by metatranscriptomics data, by conserved syntenic variants, and by previous laboratory work. Analysis of 14 metatranscriptomes (downloaded from the IMG/MER) show that multi-gene mRNAs, which likely represent operons, are present across three FaRLiP groups (Figure S5, Table S10).

Analyzing both rare and common cluster variants also suggests conserved patterns of transcription. For example, single-gene translocations and inversions have only been observed for core genes *psbA3* and the chlorophyll *f* synthase (*chlF*). This fact indicates that *chlF* and *psbA3* likely have their own promoters, which they might share with downstream genes. Other groups of genes (like *psa* genes) may share a promoter / regulatory mechanism between them, as hinted at by partial clusters (Figure S6). In addition, this study observed that most large insertions, translocations and/or split clusters were associated with the downstream area of the *rfp* genes. In order to express the rest of the FaRLiP cluster only under far-red light, the phytochrome (*rfpA*) needs to be constitutively expressed so as to detect far-red light, and the other components of the signaling cascade (*rfpB*/*C*) need to be present to transmit the signal. This has been shown by transcriptomic data and by genetic studies^33,52^. Therefore, *rpf* genes are expected to be differentially regulated to the rest of the FaRLiP cluster. A hypothesis is that a specific intergenic motif (defined as a conserved DNA sequence) controls the transcription of the FaRLiP operons, apart from *rpf* genes, under far-red light. This conserved region would likely interact with the DNA-binding residues in the regulatory protein RfpB, and potentially RfpD (Table S8) to promote transcription, and it would be expected to commonly be present upstream of *chlF* and *psbA3*.

A class of conserved, homologous, T-rich motifs was identified in FaRLiP intergenic regions (Figure 5A). These motifs were present within the first 440 bp upstream of particular genes, in all known FaRLiP lineages. BLAST searches suggested that this type of motif is specific to FaRLiP species. No known role was identified when searching motif databases with Tomtom^53^, but we hypothesize that it is likely to be regulatory. Previous work has shown that in some strains, the intergenic sequence upstream of *chlF* can drive the expression of a fluorescent reporter when induced by far-red light, in the presence of regulators RfpA/B/C^52,54^. This appears to rely on a conserved region^54^. In this study, the homologous motifs identified could be classified into three groups, from high to low conservation: *chlF* motifs, ‘long’ motifs, and ‘short’ motifs.

**Figure 5.**
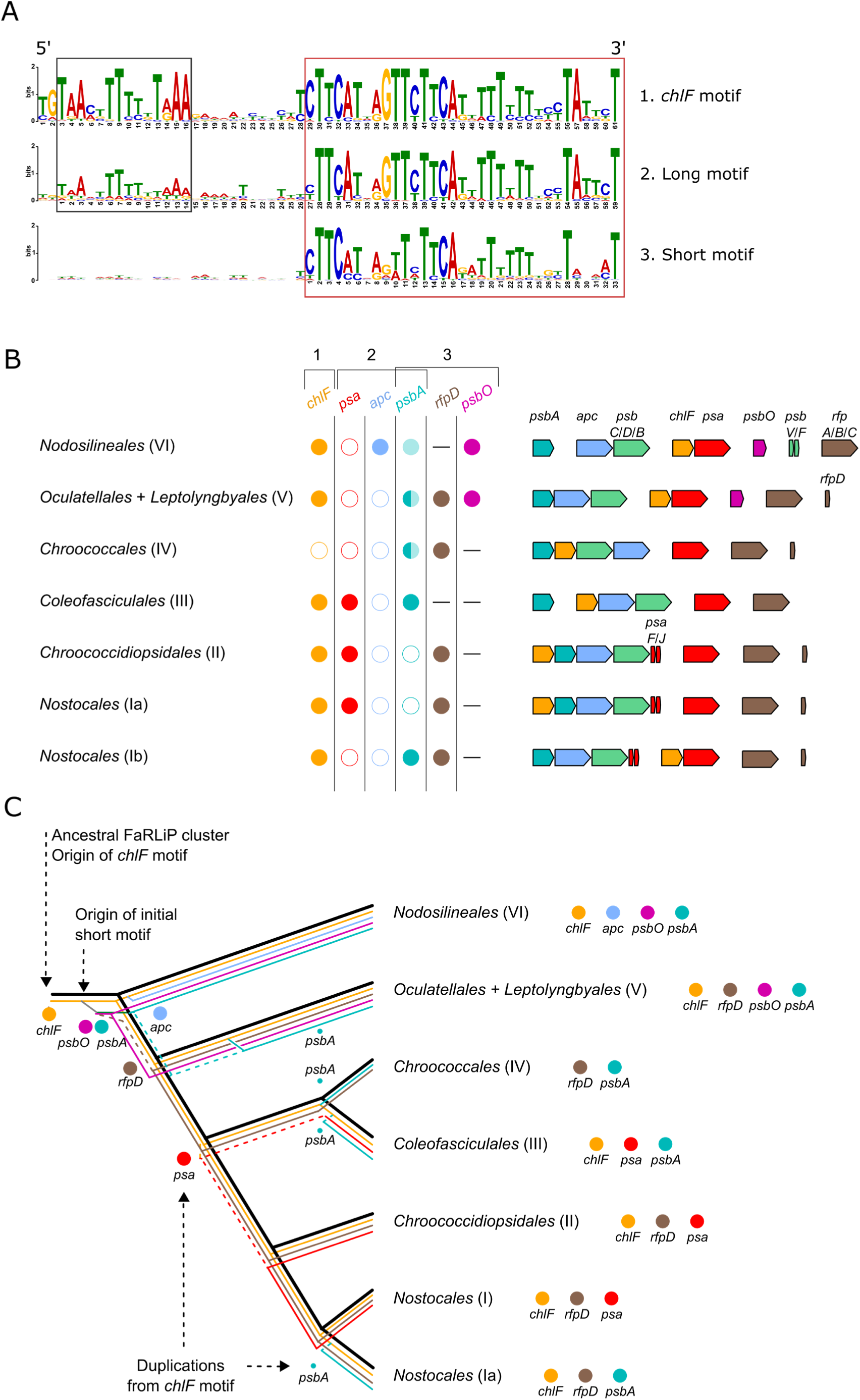
The FaRLiP gene cluster contains repeats of intergenic motifs. (A) Two distinct conserved regions, 5’ (black box) and 3’ (red box), separated by a variable gap of ∼12 bp, have been observed within them. Based on the conservation of these regions, three types of motifs (1-3) can be defined. The *chlF* motif (1) represents the most highly conserved sequence, ∼61 bp in length. It contains highly conserved 5’ and 3’ regions. Long motifs (2), ∼59 bp, are found upstream of other FaRLIP genes, but their 5’ region is less conserved. Yet other genes may only have a short motif (3), 33 bp, equivalent to only the conserved 3’ region. (B) The distribution of motifs (1-3) is shown, both according to their presence upstream of certain genes (top), as well as in particular cyanobacterial orders (left). Full circle – motif present; empty circle – motif absent, gene present; dash – gene absent. Upstream of the gene *psbA*, both long motifs (dark turquoise) and short motifs (light turquoise) may be found. Proposed operons of the FaRLiP cluster in various lineages are listed to the right. C) All FaRLiP motifs evolved from a motif preceding the gene *chlF* (yellow). From it, motifs preceding other genes were duplicated (colored lines) and their evolution is consistent with the species tree (black lines). Uncertain evolutionary paths are highlighted in dashed lines.

The most conserved motif version is present upstream of *chlF*. These *chlF* motifs are ∼61 bp long and have conserved 5’ and 3’ regions, separated by a variable gap (Figure 5A). The second group of motifs has moderate conservation, especially in the 5’ area; these are ∼59 bp ‘long’ motifs preceding other genes (*apc*, *psa*, some *psbA3* variants). The 5’ region may be lost altogether in ‘short’ motifs (∼33 bp), such as those upstream of some *psbA3* sequences, as well as minor genes *rfpD* and *psbO*. Any of these motifs is nearly always present in the intergenic region upstream of expected FaRLiP operons (Figure 5B, Table S11). Due to differences in gene arrangement, expected operons vary between lineages. Notably, no motif was detected upstream of *rfpA*/*B*/*C*, which is likely due to the different regulatory requirements of these genes.

Phylogenetic studies suggest that a highly conserved *chlF* motif was the ancestral version, from which other motifs originated through repeated, independent duplications (Figure 5C, Figure S7). These include long motifs preceding *apc* genes in the *Nodosilineales* (VI), or the *psa* genes in the *Nostocales* (I) / *Chroococcidiopsidales* (II), and some *psbA3* genes. Short motifs likely originated from a long *chlF* motif too, possibly in an early event before the split of the earliest FaRLiP lineage, the *Nodosilineales* (VI), from the rest of the cyanobacteria. All motifs, even short and divergent ones (e.g. upstream of *rfpD*), show evolutionary patterns overall supporting vertical descent. Motifs preceding *psbA3* may have a complex evolutionary history, including both long and short motifs, with ancient and recent transitions from former to latter.

The ancestral FaRLiP cluster carried a *chlF* motif. This, together with long and short motifs resulting from duplications, has been maintained across evolutionary time, some perhaps since the MRCA (2.2 – 2.5 billion years ago)^5,6^. The high conservation within the *chlF* motifs, combined with the high level of expression of the gene within the cluster^48–50^, and with the fact that this is the oldest FaRLiP gene^39,55^, underscores the vital role of the *chlf* synthase in FaRLiP. However, the co-transcription of *chlF* and *psa* genes in early lineages (V, VI) also suggests that PSI genes may have played an important role in early FaRLiP evolution.

## Discussion

Competition for light is as ancient as photosynthesis itself. Photosynthetic organisms have continuously optimized their antenna and photosystems, in order to be able to absorb and utilize the light most abundant locally. Cyanobacteria, the earliest still-extant oxygenic photosynthesizers, can acclimate to blue, green, red or far-red light by changing their pigments^17^. In plants, there exist complex regulatory mechanisms, with red/far-red phytochromes controlling shading responses, or yearly life-cycles such as flowering^56^. Far-red light is typically used for sensing, not photosynthesis, in higher plants, and is rarely used for photosynthesis in algae^57,58^. In contrast, bacteriochlorophyll-based anoxygenic photosynthesis relies on the near-infrared region, with absorption of various bacterial species ranging between 740-1050 nm^58,59^. This enables them to live not only deep in microbial mats and soil, but also around hydrothermal vents, using thermal radiation^60^. At the edge between oxygenic photosynthesis using visible light, and photosynthesis using photons too low in energy to split water, there exists far-red light photoacclimation^25,58^. FaRLiP is a remarkable process that extends the Photosynthetically Active Radiation (PAR) of oxygenic photosynthesis into the near-infrared.

This study proves that FaRLiP is an ancient phenotype. It has been inherited vertically at least since its MRCA, prior to the split of the order *Nodosilineales* (VI) from the cyanobacterial species tree. The evidence can be seen in gene trees, which tend to match the species tree, as well as in synteny, and in supporting evidence such as GC% (Table S9). Different genes were suggested to have entered the cluster at different times, with the earliest being the chl *f* synthase^39,55^. In contrast, PSI genes were suggested to have emerged later, thereby requiring HGT to explain gene presence in early lineages^38^. Our results confirm the vital role of the chl *f* synthase in the history of the cluster. However, they disagree with the HGT hypothesis. This study shows that many FaRLiP PSI genes are characterized by unusually slow/fast evolutionary rates, which make phylogenetic reconstructions difficult to build, and challenging to interpret. For now, vertical descent remains sufficient to explain the distribution of FaRLiP genes in the genomes of the lineages studied.

The genome is an ecosystem. Genes thrive or are lost depending on their environment, which is often represented by other genes. The FaRLiP cluster MRCA likely co-transcribed allophycocyanin genes with most PSII genes, while *chlF* was co-transcribed with PSI genes. The cluster may have also included a far-red *psbO*. It is not possible to tell from the synteny data whether PSI or non-*chlF* PSII genes, were the earliest. The fact that partial FaRLiP clusters exist, lacking either PSI or PSII genes, supports either option (Figure S6). The FaRLiP cluster from *Chroococcidiopsis* sp. CCMEE010, for example, lacks PSI genes, but was shown to be functional^61^. As a counterpoint, in another experiment, when the *chlF* gene was introduced into non-FaRLiP *Synechococcus* sp. PCC 7002, chl *f* was incorporated into standard PSI^62,63^. It is feasible to imagine that selective pressure in an early cyanobacterium may have led to a duplication of PSI, leading to a photosystem that could more advantageously incorporate the new chlorophyll. However, it should be noted that without a similarly red-shifted PSII, the system might not be photosynthetic (fixing carbon) but only phototropic (producing ATP). Alternatives can be seen in modern plants, which rely on both visible and far-red light photons in response to shading, and in algae, which can solely use far-red light through mechanisms yet to be fully understood^57,64,65^.

The FaRLiP cluster is characterized by regulatory simplicity. One class of highly conserved, homologous intergenic motifs may be responsible for much of its regulation. In addition, no functional FaRLiP cluster has yet to be observed to be broken into more than two pieces. Reunifications are also apparent. This stands in contrast to the complexity seen for white-light paralogues. These are commonly scattered throughout the genome and therefore transcribed and regulated independently^66,67^. Colocalization typically is limited to a few gene pairs, such as *psaA*/*B*, or *psbC*/*D*, or rarely more genes as in the phycobilisome operon *cpcB*/*A*/*C*/*C*/*D*^68,69^. A simpler and more syntenic system with fewer regulatory regions is likely to be more robust, and easier to maintain when not in use for long periods of time. Robustness has mostly been discussed for essential genes (‘the persistence model’), but recent work has extended it to genes under weak selection by considering transposable elements^70,71^.

Co-regulation appears to be a vital characteristic of the FaRLiP cluster. An additional hypothesis is that this may be connected to photosystem component replacement rates. The most easily damaged protein in white-light PSII is *psbA*^72^, whose FaRLiP version has its own promoter. Then, in order from second-most to least easily damaged, there are *psbD*, *psbC* and *psbB*^72^. It might be that by simply relying on gene order and incomplete transcripts, the FaRLiP cluster can ensure part of the regulation necessary for timely replacement. This would be consistent with transcriptional rates observed in far-red light adapted *Chlorogloeopsis fritschii* PCC 9212^49^. However, to what point transcriptional (versus translational) regulation influences the amount of end product in bacteria is still debated^73–75^.

This study was able to recover a large amount of sequence data for FaRLiP, in particular for previously underrepresented lineages, by using whole-genome metagenomics, and relying on the output of sequence search engines such as SearchSRA^76^. Other previous strategies have involved searching for strains in far-red enriched environments, or using the 16S rRNA genes of known FaRLiP strains as a proxy^28,30–32,40,77,78^. Although the genetic sequence alone does not prove chl *f* photosynthesis, the presence of a complete FaRLiP gene cluster in the present study represents strong support for it, while the recovery of partial clusters and pseudogenes was useful for uncovering patterns of evolution.

The FaRLiP cluster was likely present in the common ancestor of early-branching lineages (*Leptolyngbya*-like morphotypes, now known as *Nodosilineales*/*Oculatellales*/*Leptolyngbyales*), before their split from the rest of the cyanobacteria. In contrast with even earlier (‘basal’) branches, these cyanobacteria have been associated with evolutionary innovations in microbial mat formation, such as increased ability to tolerate salinity, a filamentous growth form and nitrogen fixation, sometimes larger cell sizes (>2.5 µm), enabling the growth of cyanobacteria in thicker layers^5–7^. The ability to photosynthesize using far-red light would likely have been beneficial in these shaded environments. Several molecular clock studies place this ancestor in the early Paleoproterozoic (or Siderian), concomitant with the rise in stromatolite diversity^4–6,12^. The presence and comparative abundance of FaRLiP species in modern microbialites, together with the presence of FaRLiP transcripts (Table S10), suggest that FaRLiP could have been highly influential in Paleoproterozoic stromatolite formation. During this time, and after the stromatolites’ decline in the present Phanerozoic eon^12^, FaRLiP cyanobacteria continued to diversify. They are present on all continents, including Antarctica, rarely in hot and cold deserts, from ocean microplastics to forest canopies and the inside of caves^26,28,31,79^, but especially in various microbial mats and modern microbialites.

## STAR Methods

### Key resource table

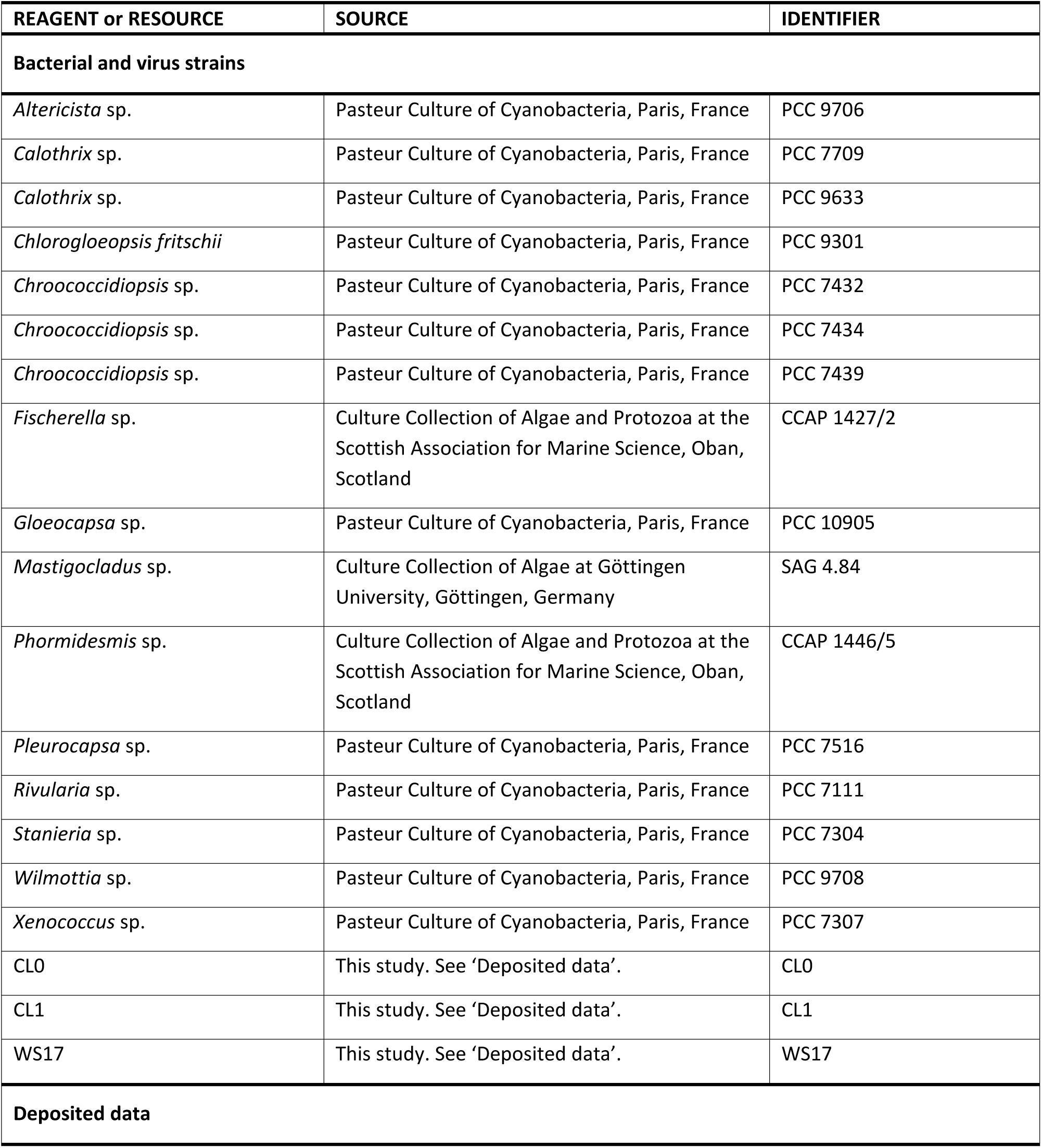
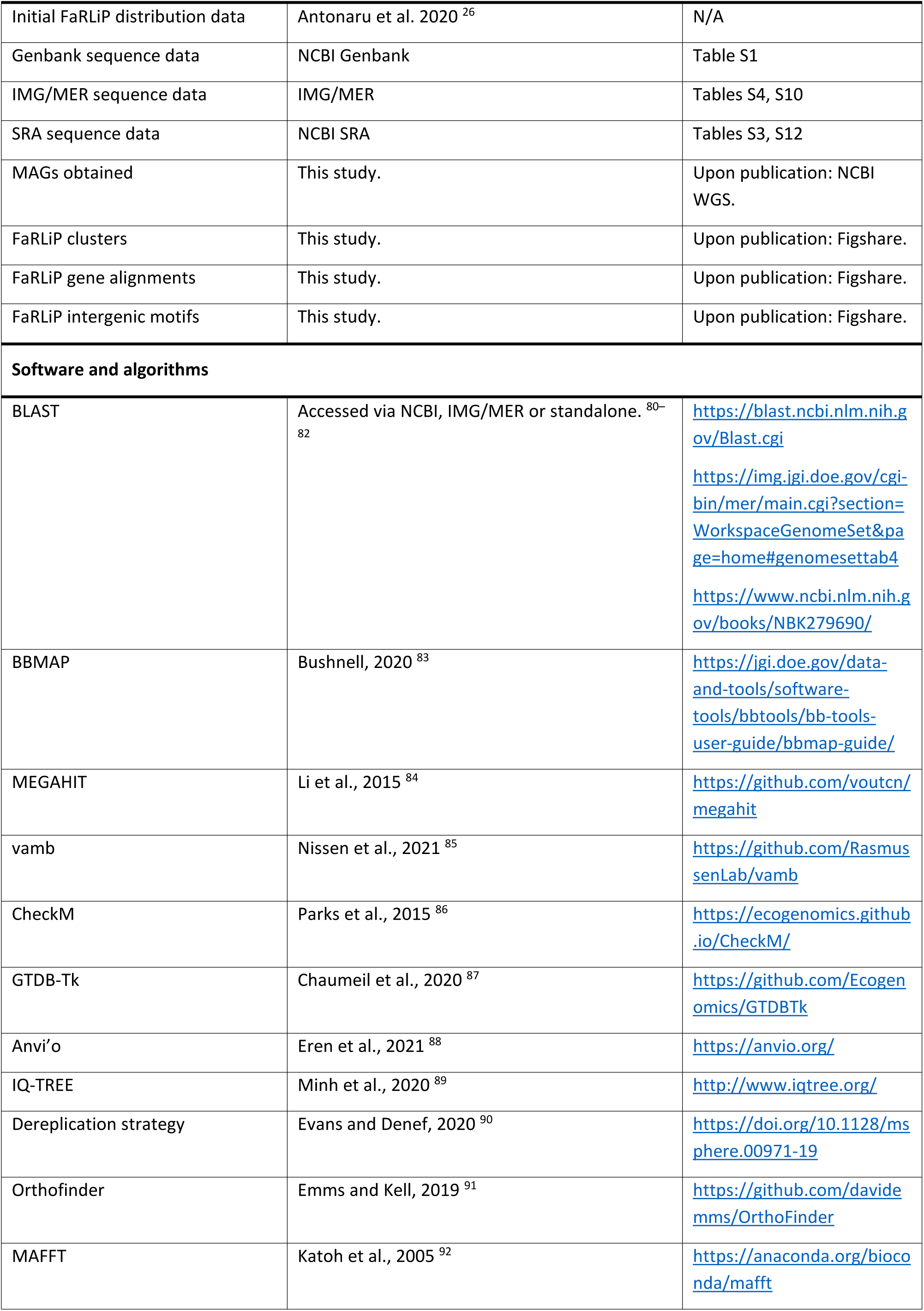
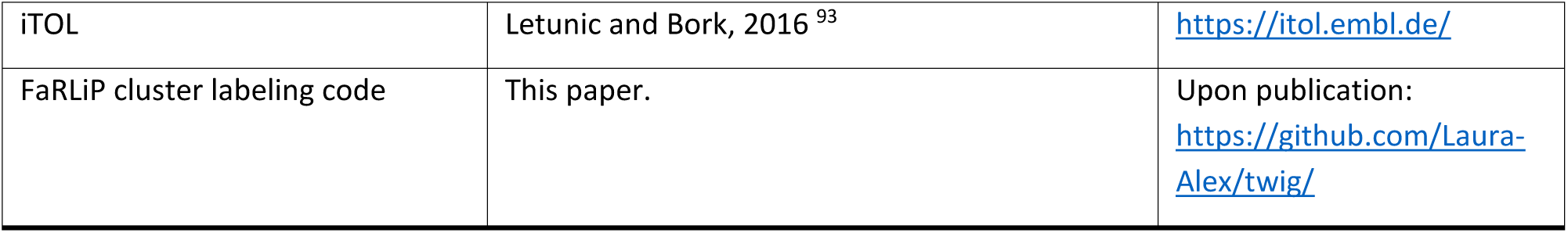

### Resource availability

#### Lead contact

Further information should be directed to and will be fulfilled by the lead contact, Dennis J. Nürnberg.

#### Materials availability

This study did not generate new unique reagents.

#### Data and code availability

This paper analyzes existing, publicly available data. These accession numbers for the datasets are listed in the Supplementary Data (see key resources table). MAGs built during the course of the study have been deposited in the NCBI WGS database and are publicly available as of the date of publication. Accession numbers are listed in Table S3. FaRLiP clusters used in this study, together with alignments of associated genes and intergenic regions, can be accessed in a figshare repository. Original code used in the cluster labeling process is accessible at <GITHUB once published>. All of these entries are also listed in the key resources table. Any additional information required to reanalyze the data reported in this paper is available from the lead contact upon request.

### Method details

#### Cyanobacterial samples

Cyanobacterial strains were used from the CCAP/SAMS (UK) and PCC (France) culture collections. Strains and culture conditions are listed in Table S2. Three additional strains were recovered from environmental samples. The CL0 and CL1 isolates were originally sampled from Lake Clifton (Australia) as previously described^26^. For WS17, salt crust fragments were sampled from the Sebkha Oum Dba, West Sahara, Morocco, in May 2019. All cultures were enriched in chlorophyll *f*-containing cyanobacteria by growing them under 730 or 750 nm light (Shenzhen Gouly LED Limited; Epitex, L750-01AU) at room temperature in liquid or on solid media (1% (w/v) agar). Strains were repeatedly re-inoculated in culture media for isolation, as previously described for CL0; for CL1 and WS17, modified DSMZ brackish medium BA+50+B+N/2 (#1678) containing a range of NaCl concentrations (0, 0.6 and 1.8% (w/v)) was used instead for the isolation, with standard BA+50+B+N/2 for later culturing. Cycloheximide (50 µg/mL) was initially added to reduce eukaryotic contamination.

#### Environmental classification

A subset of 86 FaRLiP clusters was entered into the environmental analysis. The clusters had to be either complete, near-complete (missing up to four genes in total, and up to three genes >600 bp), or (in a minority of cases) be of particular interest due to their gene rearrangements (*Nostoc* sp. C052, *Chroococcidiopsis* sp. CCMEE10). To limit the effect of sampling bias while preserving sequence diversity, virtually identical clusters from the same environment were filtered out. This includes various sequences from South African stromatolites, as well as many *Fischerella* strains from hot springs. The phylogeny of *Fischerella* strains is briefly mentioned in a previous publication^80^, with a few additional ones listed in Table S1. Identical clusters were not removed if sampled from a different environment. The working dataset was classified into four categories, as follows: microbialites, microbial mats, other (including many hot spring samples where, although the sample may have been a microbial mat, it was not specified), and unknown.

#### DNA extraction

For isolates: Genomic DNA from enriched samples was extracted using Quick-DNA Fungal/Bacterial Miniprep Kit (Zymo Research). A longer bead-beating step (25 min) was used in order to homogenize the microbial mats, together with prior fragmenting with a pipette. For CCAP strains: Cells were lysed by mechanical grinding using mortar and pestle, and genomic DNA was extracted using the DNeasy Plant Mini Kit (Qiagen), following the manufacturer instructions. For PCC strains: 40 mL of cultures in late exponential/linear growth phase were centrifuged at 12,000 x g for 10 min at 20 °C. After washing twice with sterile distilled water, or sterile saline solution (1% (w/v) NaCl) for marine strains, the pellets were immediately frozen in liquid N_2_ prior to being lyophilized. DNA of the lyophilized pellets was extracted using NucleoBond AXG 20 columns (Macherey-Nagel) according to the manufacturer’s instructions for bacterial DNA.

#### (Meta)genome sequencing

For isolates: Sample CL0 was sequenced with both Illumina and Oxford Nanopore as previously described for other strains^79^. It was sequenced according to vendor protocols on a MiSeq and MinION (MinKNOW version 1.10.23), the former using Nextera XT v2 and the latter a R9.4.1 flowcell with base-calling via Albacore (version 2.1.7). Samples CL1 and WS17 were sequenced using Illumina. For these two samples, library preparation was done using NEBNext Ultra II DNA Library Prep Kit (New England Biolabs) in combination with NEBNext Multiplex Oligos for Illumina (Unique Dual Index UMI Adaptors) according to the manufacturer’s manual. The libraries were sequenced on Illumina MiSeq using the sequencing kit v3 600 cycles (PE 2×300). For CCAP strains: Paired-end libraries of 150 bp (PE150) were built from genomic DNA, and sequenced using Illumina NovaSeq 6000 to a depth of ca. 40 million raw reads (ca. 6 Gb raw data) per library (Novogene Ltd). For PCC strains: The genomes were sequenced with Illumina NextSeq500 system from the Mutualized Platform for Microbiology (P2M) of the Institut Pasteur or from GATC.

#### (Meta)genome assembly

For isolates and datasets from sequence databases: The metagenomes were quality-trimmed with BBDuk (software package BBMap) ^81^. Settings for Illumina: qtrim=r trimq=10 minlen=30 ref=adapters.fa (default) ktrim=r k=23 mink=11 hdist=1 tpe tbo. Settings for Nanopore: trimq=9 ktrim=l k=11 hdist=0 edist=2 mm=f rcomp=f mkf=0.51 restrictleft=100 copyundefined=t. The results were assembled with MEGAHIT^82^, with the exception of less-diverse CL1 and WS17 samples which used SPADES^83^, and the sample CL0 which used Unicycler^84^ to incorporate the long reads. Paired-end information was used if available. For the cluster shape analysis, sequences in the same environment were co-assembled where computationally possible (Table S12), while for building MAGs, they were assembled individually and then co-binned (Table S3). The latter is expected to be more accurate, while the former led to more complete FaRLiP clusters. The two methods agreed on the sequence level. For CCAP strains: A metagenomic pipeline based on metaWRAP^85^ was used to pre-process sequence data for quality and assemble using metaSPADES (v3.13.0)^86^. For PCC strains: Assemblies were built using fq2dna v21.06^87^, with coverage ranging from 58x to 438x. PCC cluster sequences are available under accession numbers: ((will be provided)).

#### Metagenome binning and quality control

For CCAP strains, ensemble metagenome binning was based on Concoct^88^, MaxBin2^89^ and MetaBat2^90^. For isolates and datasets from sequence databases, a different method was used. Independently assembled samples from the same environment were co-binned with vamb^91^. Bin quality was checked with CheckM^92^. Bins with >70% completion were further manually refined with anvi’o^93^, and the final refined dataset aimed for bins with >70% completion and <5% redundancy, also based on CheckM (Table S3).

Duplicate bins from the same environments were removed with MASH and pyANI, using a de-replication strategy and code previously described ^94^. Remaining bins which were cyanobacterial in nature and contained FaRLiP genes were used in the study. A sequence similarity analysis was performed for groups V and VI with EzAAI at both genome and cluster level, in order to better understand a divergent group V lineage^95^. In this case, the cluster was approximated as the concatenation of 19 core genes.

For all assemblies, the binned metagenome assembled genomes were verified using CheckM^92^, taxonomically identified with GTDB-tk^96^ and annotated using Prokka^97^. All bins used in this study are available at ((NCBI codes will be provided)).

#### Database sourcing

The main databases used in this study: NCBI (nr, WGS and SRA) and IMG/MER^98–100^. For the NCBI nr (non-redundant) and WGS (Whole Genome Shotgun) databases, BLASTp and tBLASTn were used to recover new genes and clusters ^101,102^. The far-red specific ApcE2 motif VIPEDV was used as a query, as previously described^26^. IMG/MER and SRA data is based on accessions originally recovered through BLAST or SearchSRA^76^ in previous work with the same query^26^. The IMG dataset also contained metatranscriptomics. SRA data included all runs from the projects previously identified to contain a VIPEDV-like motif^26^, which were then downloaded with fasterq-dump^103^.

Other FaRLiP genes besides *apcE2* were recovered largely via co-localization with *apcE2*. However, BLASTp searches of other FaRLiP genes and motifs (Table S3) were also conducted, in order to test that the previously recovered genes were indeed far-red paralogues, and to recover split clusters, partial clusters and pseudogenes. This was applied to fragmented data, either from public databases or locally assembled. Gene alignments were constructed with MAFFT in Jalview (default settings)^104,105^. SplitsTree 5 software^106^ was used to build networks of the aligned genes and to rapidly identify sequences clustering with known FaRLiP paralogues.

#### Phylogenetic analysis

DNA and protein alignments for the 19 core genes (present in all complete FaRLiP clusters) were constructed with MAFFT (--maxiterate 1000 --localpair). Alignments included 81-96 sequences each, depending on the gene. General sequence management such as renaming was performed with SeqKit^107^. Trees were inferred with IQ-TREE (bootstrap 1000), with the best substitution model being chosen automatically using ModelFinder^108^. As per the HoT method, both ‘head’ (default) and ‘tails’ (reverse-complement, align, then reverse-complement again) alignments and trees were built, for both protein and DNA sequences, in order to assess alignment reliability^47^. To further test possible effects of alignment artifacts, head and tails trees were compared with IQ-TREE’s –sup argument. Trees were rooted with Minimal Ancestral Deviation (MAD)^109^, with the best-scoring root point (lowest Ambiguity Index in multiple trees) supporting vertical descent (Table S13). Trees were illustrated with iTOL and Inkscape 1.2.1^110^. SplitsTree 5 was used to group multiple trees into networks^106^.

A calculation of synonymous and non-synonymous substitution rates (dS, dN) was executed with a combination of MAFFT (ClustalW format), pal2nal (aligning protein to nucleotide sequences) and codeml (rate calculation)^111,112^.

The genome tree was built with Orthofinder^113^ and is consistent with recent work^13^. It focused on FaRLiP genomes and MAGs, but also included entries lacking the cluster (*Gloeobacter violaceous* PCC 7421, *‘Chroococcidiopsis*’ sp. CCMEE 29) or containing only some FaRLiP genes (*Hyella patelloides* LEGE 07179) for increased phylogenetic accuracy.

#### Synteny analysis

Annotated, manually corrected clusters were submitted to the Gene Graphics webserver for visualization^114^, and then compared for similarities. This required a change in data formatting (from the .gbk Prokka output, to a .tsv tabular format for Gene Graphics). The code used can be found at <GITHUB once published>. Final figures created with Inkscape 1.2.1.

#### Taxonomic conventions

This paper uses a genome-based taxonomy^13^. For ease of cross-referencing with the GTDB classification, or NCBI taxonomy, see Table S14.

#### Motif identification

The *chlF* motif and several long motifs were identified by aligning intergenic areas upstream of their respective genes with MAFFT, in Jalview. Less-conserved motifs were recovered by aligning the remaining intergenic areas with multiple copies of the conserved *chlF* motifs, thus forcing alignment to this region. Results were checked with the MEME webserver, which was also used to obtain the consensus logos. MEME cannot use gapped data, and so occasional single-nucleotide deletions were replaced with “X” in the dataset.

## Acknowledgements

Many thanks to Tal Dagan and to Michael W. Gaunt for bioinformatics advice. DJN acknowledges the support of Europlanet 2020 RI for sampling in Morocco. Europlanet 2020 RI has received funding from the European Union’s Horizon 2020 research and innovation programme under grant agreement No 654208. The HPC facilities at Imperial College London, Freie Universität Berlin and University of Kiel were used during the study. Daniel J. Wilson, Nicholas D. Sanderson and Leanne Barker are thanked for the sequencing of the CL0 and SAG 4.84 clusters, with sequencing supported by the National Institute for Health Research Health Protection Research Unit (NIHR HPRU) at the University of Oxford in partnership with Public Health England (PHE) (HPRU-2012-10041 to DJW). This research was supported by the Deutsche Forschungsgemeinschaft (DFG; Emmy Noether project award no. NU 421/1 to DJN). LAA was supported by a postdoctoral fellowship from the Freie Universität Berlin and a Schrödinger fellowship from Imperial College London. SW by a scholarship from the China Scholarship Council. Access to the CCAP/SAMS collection was possible through the AssemblePlus program. CRM and DG were funded in part by the UK Natural Environment research Centre (project NE/R017050/1). The Pasteur Cultures of Cyanobacteria is funded by the Institut Pasteur. The Mutualized Platform for Microbiology (P2M) is acknowledged for PCC genomes. Open access funding provided by Project DEAL.

## Author contributions

L.A.A. and D.J.N. designed the study. C.R-M., D.G. and M.G. provided strains and sequence information. S.W. cultured the strains. S.M., S.S. and D.G. were responsible for sequencing and data curation. L.A.A. performed the analyses with the help of M.P. and T.O. for the intergenic motif analysis. L.A.A. and D.J.N. wrote the paper, with contributions from all authors.

## Supplemental information

Supplementary information file. Figures S1-S7, and tables S2-S10, S12-S14.

Table S1+S11. Tables S1 (NCBI data) and S11 (Intergenic motifs) are provided as an excel file.

## References

1. Lyons, T.W., Reinhard, C.T., and Planavsky, N.J. (2014). The rise of oxygen in Earth’s early ocean and atmosphere. Nature 506, 307–315. 10.1038/nature13068.

2. Schopf, J.W. (2011). The paleobiological record of photosynthesis. Photosynth. Res. 107, 87–101. 10.1007/s11120-010-9577-1.

3. Flombaum, P., Gallegos, J.L., Gordillo, R.A., Rincón, J., Zabala, L.L., Jiao, N., Karl, D.M., Li, W.K.W., Lomas, M.W., Veneziano, D., et al. (2013). Present and future global distributions of the marine Cyanobacteria Prochlorococcus and Synechococcus. Proc. Natl. Acad. Sci. U. S. A. 110, 9824–9829. 10.1073/pnas.1307701110.

4. Knoll, A.H. (2008). Cyanobacteria and Earth History. In The Cyanobacteria: Molecular Biology, Genomics and Evolution, A. Herrero and E. Flores, eds. (Caister Academic Press), pp. 1–20.

5. Blank, C.E., and Sanchez-Baracaldo, P. (2010). Timing of morphological and ecological innovations in the cyanobacteria - A key to understanding the rise in atmospheric oxygen. Geobiology 8, 1–23. 10.1111/j.1472-4669.2009.00220.x.

6. Uyeda, J.C., Harmon, L.J., and Blank, C.E. (2016). A comprehensive study of cyanobacterial morphological and ecological evolutionary dynamics through deep geologic time. PLoS One 11, 1–32. 10.1371/journal.pone.0162539.

7. Schirrmeister, B.E., Gugger, M., and Donoghue, P.C.J. (2015). Cyanobacteria and the Great Oxidation Event: Evidence from genes and fossils. Palaeontology 58, 769–785. 10.1111/pala.12178.

8. Demoulin, C.F., Lara, Y.J., Lambion, A., and Javaux, E.J. (2024). Oldest thylakoids in fossil cells directly evidence oxygenic photosynthesis. Nature 625, 529–534. 10.1038/s41586-023-06896-7.

9. Vítek, P., Ascaso, C., Artieda, O., Casero, M.C., and Wierzchos, J. (2017). Discovery of carotenoid red-shift in endolithic cyanobacteria from the Atacama Desert. Sci. Rep. 7, 1–10. 10.1038/s41598-017-11581-7.

10. Schirrmeister, B.E., Antonelli, A., and Bagheri, H.C. (2011). The origin of multicellularity in cyanobacteria. BMC Evol. Biol. 11, 1–21. 10.1186/1471-2148-11-45.

11. Boden, A.J.S., Nieves-Morión, M., Nürnberg, D.J., Arévalo, S., Flores, E., and Sánchez-Baracaldo, P. (2023). Evolution of Multicellularity Genes in the Lead Up to the Great Oxidation Event. bioRxiv December, 1–33. 10.1101/2023.12.23.573081.

12. Awramik, S.M., and Sprinkle, J. (1999). Proterozoic stromatolites: The first Marine Evolutionary Biota. Hist. Biol. 13, 241–253. 10.1080/08912969909386584.

13. Strunecký, O., Ivanova, A.P., and Mareš, J. (2022). An updated classification of cyanobacterial orders and families based on phylogenomic and polyphasic analysis. J. Phycol. 51, 12–51. 10.1111/jpy.13304.

14. Fiebig, O.C., Harris, D., Wang, D., Hoffmann, M.P., and Schlau-Cohen, G.S. (2023). Ultrafast Dynamics of Photosynthetic Light Harvesting: Strategies for Acclimation Across Organisms. Annu. Rev. Phys. Chem. 74, 493–520. 10.1146/annurev-physchem-083122-111318.

15. Holtrop, T., Huisman, J., Stomp, M., Biersteker, L., Aerts, J., Grébert, T., Partensky, F., Garczarek, L., and van der Woerd, H.J. (2020). Vibrational modes of water predict spectral niches for photosynthesis in lakes and oceans. Nat. Ecol. Evol. 5, 55–66. 10.1038/s41559-020-01330-x.

16. Gan, F., and Bryant, D.A. (2015). Adaptive and acclimative responses of cyanobacteria to far-red light. Environ. Microbiol. 17, 3450–3465. 10.1111/1462-2920.12992.

17. Sanfilippo, J.E., Garczarek, L., Partensky, F., and Kehoe, D.M. (2019). Chromatic acclimation in cyanobacteria: A diverse and widespread process for optimizing photosynthesis. Annu. Rev. Microbiol. 73, 407–433. 10.1146/annurev-micro-020518-115738.

18. Stomp, M., Huisman, J., Stal, L.J., and Matthijs, H.C.P. (2007). Colorful niches of phototrophic microorganisms shaped by vibrations of the water molecule. ISME J. 1, 271–282. 10.1038/ismej.2007.59.

19. Grébert, T., Doré, H., Partensky, F., Farrant, G.K., Boss, E.S., Picheral, M., Guidi, L., Pesant, S., Scanlan, D.J., Wincker, P., et al. (2018). Light color acclimation is a key process in the global ocean distribution of Synechococcus cyanobacteria. Proc. Natl. Acad. Sci. U. S. A. 115, E2010–E2019. 10.1073/pnas.1717069115.

20. Chen, M.Y., Teng, W.K., Zhao, L., Hu, C.X., Zhou, Y.K., Han, B.P., Song, L.R., and Shu, W.S. (2020). Comparative genomics reveals insights into cyanobacterial evolution and habitat adaptation. ISME J. 15, 211–227. 10.1038/s41396-020-00775-z.

21. Wang, F., and Chen, M. (2022). Chromatic Acclimation Processes and Their Relationships with Phycobiliprotein Complexes. Microorganisms 10. 10.3390/microorganisms10081562.

22. Nürnberg, D.J., Morton, J., Santabarbara, S., Telfer, A., Joliot, P., Antonaru, L.A., Ruban, A. V., Cardona, T., Krausz, E., Boussac, A., et al. (2018). Photochemistry beyond the red limit in chlorophyll f–containing photosystems. Science 360, 1210–1213. 10.1126/science.aar8313.

23. Chen, M., Li, Y., Birch, D., and Willows, R.D. (2012). A cyanobacterium that contains chlorophyll f - A red-absorbing photopigment. FEBS Lett. 586, 3249–3254. 10.1016/j.febslet.2012.06.045.

24. Gan, F., Zhang, S., Rockwell, N.C., Martin, S.S., Lagarias, J.C., and Bryant, D.A. (2014). Extensive remodeling of a cyanobacterial photosynthetic apparatus in far-red light. Science 345, 1312–1317. 10.1126/science.1256963.

25. Elias, E., Oliver, T.J., and Croce, R. (2024). Oxygenic Photosynthesis in Far-Red Light: Strategies and Mechanisms. Annu. Rev. Phys. Chem. 75, 231–256.

26. Antonaru, L.A., Cardona, T., Larkum, A.W.D., and Nürnberg, D.J. (2020). Global distribution of a chlorophyll f cyanobacterial marker. ISME J. 14, 2275–2287. 10.1038/s41396-020-0670-y.

27. Averina, S., Velichko, N., Senatskaya, E., and Pinevich, A. (2018). Far-red light photoadaptations in aquatic cyanobacteria. Hydrobiologia 813, 1–17. 10.1007/s10750-018-3519-x.

28. Zhang, Z.C., Li, Z.K., Yin, Y.C., Li, Y., Jia, Y., Chen, M., and Qiu, B.S. (2019). Widespread occurrence and unexpected diversity of red-shifted chlorophyll producing cyanobacteria in humid subtropical forest ecosystems. Environ. Microbiol. 21, 1497–1510. 10.1111/1462-2920.14582.

29. Gómez-Lojero, C., Leyva-Castillo, L.E., Herrera-Salgado, P., Barrera-Rojas, J., Ríos-Castro, E., and Gutiérrez-Cirlos, E.B. (2018). Leptolyngbya CCM 4, a cyanobacterium with far-red photoacclimation from Cuatro Ciénegas Basin, México. Photosynthetica 56, 342–353. 10.1007/s11099-018-0774-z.

30. Kühl, M., Trampe, E., Mosshammer, M., Johnson, M., Larkum, A.W.D., and Koren, K. (2020). Substantial near-infrared radiation-driven photosynthesis of chlorophyll f-containing cyanobacteria in a natural habitat. Elife 1, 1–15. 10.1101/750174.

31. Behrendt, L., Trampe, E.L., Nord, N.B., Nguyen, J., Kühl, M., Lonco, D., Nyarko, A., Dhinojwala, A., Hershey, O.S., and Barton, H. (2020). Life in the dark: far-red absorbing cyanobacteria extend photic zones deep into terrestrial caves. Environ. Microbiol. 22, 952–963. 10.1111/1462-2920.14774.

32. Ohkubo, S., and Miyashita, H. (2017). A niche for cyanobacteria producing chlorophyll f within a microbial mat. ISME J. 11, 2368–2378. 10.1038/ismej.2017.98.

33. Zhao, C., Gan, F., Shen, G., and Bryant, D.A. (2015). RfpA, RfpB, and RfpC are the master control elements of far-red light photoacclimation (FaRLiP). Front. Microbiol. 6, 1–13. 10.3389/fmicb.2015.01303.

34. Ho, M.Y., Shen, G., Canniffe, D.P., Zhao, C., and Bryant, D.A. (2016). Light-dependent chlorophyll f synthase is a highly divergent paralog of PsbA of photosystem II. Science 353, 1–7. 10.1126/science.aaf9178.

35. Soulier, N., Laremore, T.N., and Bryant, D.A. (2020). Characterization of cyanobacterial allophycocyanins absorbing far-red light. Photosynth. Res. 145, 189–207. 10.1007/s11120-020-00775-2.

36. Trinugroho, J.P., Bečková, M., Shao, S., Yu, J., Zhao, Z., Murray, J.W., Sobotka, R., Komenda, J., and Nixon, P.J. (2020). Chlorophyll f synthesis by a super-rogue photosystem II complex. Nat. Plants 6, 238–244. 10.1038/s41477-020-0616-4.

37. Gan, F., Shen, G., and Bryant, D.A. (2015). Occurrence of far-red light photoacclimation (FaRLiP) in diverse cyanobacteria. Life 5, 4–24. 10.3390/life5010004.

38. Gisriel, C.J., Bryant, D.A., Brudvig, G.W., and Cardona, T. (2023). Molecular diversity and evolution of far-red light-acclimated photosystem I. Front. Plant Sci. 14, 1–16. 10.3389/fpls.2023.1289199.

39. Gisriel, C.J., Cardona, T., Bryant, D.A., and Brudvig, G.W. (2022). Molecular Evolution of Far-Red Light-Acclimated Photosystem II. Microorganisms 10, 1–18. 10.3390/microorganisms10071270.

40. Ko, J.T., Li, Y.Y., Chen, P.Y., Liu, P.Y., and Ho, M.Y. (2023). Use of 16S rRNA gene sequences to identify cyanobacteria that can grow in far-red light. Mol. Ecol. Resour. 24, 1–15. 10.1111/1755-0998.13871.

41. Oliver, T., Kim, T.D., Trinugroho, J.P., Cordón-Preciado, V., Wijayatilake, N., Bhatia, A., Rutherford, A.W., and Cardona, T. (2023). The Evolution and Evolvability of Photosystem II. Annu. Rev. Plant Biol. 74, 225–257. 10.1146/annurev-arplant-070522-062509.

42. Averina, S., Polyakova, E., Senatskaya, E., and Pinevich, A. (2021). A new cyanobacterial genus Altericista and three species, A. lacusladogae sp. nov., A. violacea sp. nov., and A. variichlora sp. nov., described using a polyphasic approach. J. Phycol. 57, 1517–1529. 10.1111/jpy.13188.

43. Sheridan, K.J., Duncan, E.J., Eaton-Rye, J.J., and Summerfield, T.C. (2020). The diversity and distribution of D1 proteins in cyanobacteria. Photosynth. Res. 145, 111–128. 10.1007/s11120-020-00762-7.

44. Bryant, J.A., Clemente, T.M., Viviani, D.A., Fong, A.A., Thomas, K.A., Kemp, P., Karl, D.M., White, A.E., and DeLong, E.F. (2016). Diversity and Activity of Communities Inhabiting Plastic Debris in the North Pacific Gyre. mSystems 1, 1–19. 10.1128/msystems.00024-16.

45. Zettler, E.R., Mincer, T.J., and Amaral-Zettler, L.A. (2013). Life in the “plastisphere”: Microbial communities on plastic marine debris. Environ. Sci. Technol. 47, 7137–7146. 10.1021/es401288x.

46. Philippe, H., Brinkmann, H., Lavrov, D. V., Littlewood, D.T.J., Manuel, M., Wörheide, G., and Baurain, D. (2011). Resolving difficult phylogenetic questions: Why more sequences are not enough. PLoS Biol. 9, 1–10. 10.1371/journal.pbio.1000602.

47. Landan, G., and Graur, D. (2007). Heads or tails: A simple reliability check for multiple sequence alignments. Mol. Biol. Evol. 24, 1380–1383. 10.1093/molbev/msm060.

48. Chen, M., Hernandez-Prieto, M.A., Loughlin, P.C., Li, Y., and Willows, R.D. (2019). Genome and proteome of the chlorophyll f-producing cyanobacterium Halomicronema hongdechloris: Adaptative proteomic shifts under different light conditions. BMC Genomics 20, 1–15. 10.1186/s12864-019-5587-3.

49. Ho, M.Y., and Bryant, D.A. (2019). Global transcriptional profiling of the cyanobacterium Chlorogloeopsis fritschii PCC 9212 in far-red light: Insights into the regulation of chlorophyll d synthesis. Front. Microbiol. 10, 1–16. 10.3389/fmicb.2019.00465.

50. Ho, M.Y., Gan, F., Shen, G., Zhao, C., and Bryant, D.A. (2017). Far-red light photoacclimation (FaRLiP) in Synechococcus sp. PCC 7335: I. Regulation of FaRLiP gene expression. Photosynth. Res. 131, 173–186. 10.1007/s11120-016-0309-z.

51. Gisriel, C.J., Shen, G., Ho, M.-Y.Y., Kurashov, V., Flesher, D.A., Wang, J., Armstrong, W.H., Golbeck, J.H., Gunner, M.R., Vinyard, D.J., et al. (2022). Structure of a monomeric photosystem II core complex from a cyanobacterium acclimated to far-red light reveals the functions of chlorophylls d and f. J. Biol. Chem. 298, 1–15. 10.1016/j.jbc.2021.101424.

52. Liu, T.S., Wu, K.F., Jiang, H.W., Chen, K.W., Nien, T.S., Bryant, D.A., and Ho, M.Y. (2023). Identification of a Far-Red Light-Inducible Promoter that Exhibits Light Intensity Dependency and Reversibility in a Cyanobacterium. ACS Synth. Biol. 12, 1320–1330. 10.1021/acssynbio.3c00066.

53. Gupta, S., Stamatoyannopoulos, J.A., Bailey, T.L., and Noble, W.S. (2007). Quantifying similarity between motifs. Genome Biol. 8, R24. 10.1186/gb-2007-8-2-r24.

54. Pope, M. (2021). Improving Synechocystis sp. PCC 6803 as a model organism. PhD thesis. Imperial College London, UK.

55. Oliver, T., Sánchez-Baracaldo, P., Larkum, A.W., Rutherford, A.W., and Cardona, T. (2021). Time-resolved comparative molecular evolution of oxygenic photosynthesis. Biochim. Biophys. Acta - Bioenerg. 1862, 1–49. 10.1016/j.bbabio.2021.148400.

56. Demotes-Mainard, S., Péron, T., Corot, A., Bertheloot, J., Le Gourrierec, J., Pelleschi-Travier, S., Crespel, L., Morel, P., Huché-Thélier, L., Boumaza, R., et al. (2016). Plant responses to red and far-red lights, applications in horticulture. Environ. Exp. Bot. 121, 4–21. 10.1016/j.envexpbot.2015.05.010.

57. Wolf, B.M., and Blankenship, R.E. (2019). Far-red light acclimation in diverse oxygenic photosynthetic organisms. Photosynth. Res. 142, 349–359. 10.1007/s11120-019-00653-6.

58. Larkum, A.W.D., Ritchie, R.J., and Raven, J.A. (2018). Living off the Sun: chlorophylls, bacteriochlorophylls and rhodopsins. Photosynthetica 56, 11–43. 10.1007/s11099-018-0792-x.

59. George, D.M., Vincent, A.S., and Mackey, H.R. (2020). An overview of anoxygenic phototrophic bacteria and their applications in environmental biotechnology for sustainable Resource recovery. Biotechnol. Reports 28, e00563. 10.1016/j.btre.2020.e00563.

60. Beatty, J.T., Overmann, J., Lince, M.T., Manske, A.K., Lang, A.S., Blankenship, R.E., Van Dover, C.L., Martinson, T.A., and Plumley, F.G. (2005). An obligately photosynthetic bacterial anaerobe from a deep-sea hydrothermal vent. Proc. Natl. Acad. Sci. U. S. A. 102, 9306–9310. 10.1073/pnas.0503674102.

61. Billi, D., Napoli, A., Mosca, C., Fagliarone, C., Carolis, R. De, Balbi, A., Scanu, M., Selinger, V.M., Antonaru, L.A., and Nürnberg, D.J. (2022). Identification of far-red light acclimation in an endolithic Chroococcidiopsis strain and associated genomic features : Implications for oxygenic photosynthesis on exoplanets. Front. Microbiol. 13, 1–12. 10.3389/fmicb.2022.933404.

62. Tros, M., Bersanini, L., Shen, G., Ho, M.Y., van Stokkum, I.H.M., Bryant, D.A., and Croce, R. (2020). Harvesting far-red light: Functional integration of chlorophyll f into Photosystem I complexes of Synechococcus sp. PCC 7002. Biochim. Biophys. Acta - Bioenerg. 1861, 148206. 10.1016/j.bbabio.2020.148206.

63. Shen, G., Canniffe, D.P., Ho, M.Y., Kurashov, V., van der Est, A., Golbeck, J.H., and Bryant, D.A. (2019). Characterization of chlorophyll f synthase heterologously produced in Synechococcus sp. PCC 7002. Photosynth. Res. 140, 77–92. 10.1007/s11120-018-00610-9.

64. Melis, A., and Harvey, G.W. (1981). Regulation of photosystem stoichiometry, chlorophyll a and chlorophyll b content and relation to chloroplast ultrastructure. Biochim. Biophys. Acta 637, 138–145.

65. Kosugi, M., Kawasaki, M., Shibata, Y., Hara, K., Takaichi, S., Moriya, T., Adachi, N., Kamei, Y., Kashino, Y., Kudoh, S., et al. (2023). Uphill energy transfer mechanism for photosynthesis in an Antarctic alga. Nat. Commun. 14, 1–14. 10.1038/s41467-023-36245-1.

66. Lindell, D., Sullivan, M.B., Johnson, Z.I., Tolonen, A.C., Rohwer, F., and Chisholm, S.W. (2004). Transfer of photosynthesis genes to and from Prochlorococcus viruses. Proc. Natl. Acad. Sci. U. S. A. 101, 11013–11018. 10.1073/pnas.0401526101.

67. Korbel, J.O., Jensen, L.J., Von Mering, C., and Bork, P. (2004). Analysis of genomic context: Prediction of functional associations from conserved bidirectionally transcribed gene pairs. Nat. Biotechnol. 22, 911–917. 10.1038/nbt988.

68. Adachi, Y., Kuroda, H., Yukawa, Y., and Sugiura, M. (2012). Translation of partially overlapping psbD-psbC mRNAs in chloroplasts: The role of 5′-processing and translational coupling. Nucleic Acids Res. 40, 3152–3158. 10.1093/nar/gkr1185.

69. Shi, T., Bibby, T.S., Jiang, L., Irwin, A.J., and Falkowski, P.G. (2005). Protein interactions limit the rate of evolution of photosynthetic genes in cyanobacteria. Mol. Biol. Evol. 22, 2179–2189. 10.1093/molbev/msi216.

70. Fang, G., Rocha, E.P.C., and Danchin, A. (2008). Persistence drives gene clustering in bacterial genomes. BMC Genomics 9, 1–14. 10.1186/1471-2164-9-4.

71. Kanai, Y., Tsuru, S., and Furusawa, C. (2022). Experimental demonstration of operon formation catalyzed by insertion sequence. Nucleic Acids Res. 50, 1673–1686. 10.1093/nar/gkac004.

72. Yao, D.C.I., Brune, D.C., and Vermaas, W.F.J. (2012). Lifetimes of photosystem i and II proteins in the cyanobacterium Synechocystis sp. PCC 6803. FEBS Lett. 586, 169–173. 10.1016/j.febslet.2011.12.010.

73. Picard, F., Milhem, H., Loubière, P., Laurent, B., Cocaign-Bousquet, M., and Girbal, L. (2012). Bacterial translational regulations: High diversity between all mRNAs and major role in gene expression. BMC Genomics 13. 10.1186/1471-2164-13-528.

74. Favate, J.S., Liang, S., Cope, A.L., Yadavalli, S.S., and Shah, P. (2022). The landscape of transcriptional and translational changes over 22 years of bacterial adaptation. Elife 11, 1–25. 10.7554/eLife.81979.

75. Li, G.W., Burkhardt, D., Gross, C., and Weissman, J.S. (2014). Quantifying absolute protein synthesis rates reveals principles underlying allocation of cellular resources. Cell 157, 624–635. 10.1016/j.cell.2014.02.033.

76. Levi, K., Abeysinghe, E., Rynge, M., and Edwards, R.A. (2018). Searching the sequence read archive using Jetstream and Wrangler. In Practice and Experience in Advanced Research Computing ‘18 (PEARC ‘18), pp. 1–7. 10.1145/3219104.3229278.

77. Shen, L.Q., Zhang, Z.C., Shang, J.L., Li, Z.K., Chen, M., Li, R., and Qiu, B.S. (2022). Kovacikia minuta sp. nov. (Leptolyngbyaceae, Cyanobacteria), a new freshwater chlorophyll f-producing cyanobacterium. J. Phycol. 58, 424–435. 10.1111/jpy.13248.

78. Murray, B., Ertekin, E., Dailey, M., Soulier, N.T., Shen, G., Bryant, D.A., Perez-Fernandez, C., and Diruggiero, J. (2022). Adaptation of Cyanobacteria to the endolithic light spectrum in hyper-arid deserts. Microorganisms 10, 1–12. 10.3390/microorganisms10061198.

79. Antonaru, L.A., Selinger, V.M., Jung, P., Di Stefano, G., Sanderson, N.D., Barker, L., Wilson, D.J., Büdel, B., Canniffe, D.P., Billi, D., et al. (2023). Common loss of far-red light photoacclimation in cyanobacteria from hot and cold deserts: a case study in the Chroococcidiopsidales. ISME Commun. 3, 1–9. 10.1038/s43705-023-00319-4.

80. Alcorta, J., Vergara-Barros, P., Antonaru, L.A., Alcamán-Arias, M.E., Nürnberg, D.J., and Díez, B. (2019). Fischerella thermalis: a model organism to study thermophilic diazotrophy, photosynthesis and multicellularity in cyanobacteria. Extremophiles 23, 635–647. 10.1007/s00792-019-01125-4.

81. Bushnell, B. (2020). BBMap. BBMap short read aligner, other bioinformatic tools. https://sourceforge.net/projects/bbmap/.

82. Li, D., Liu, C.M., Luo, R., Sadakane, K., and Lam, T.W. (2015). MEGAHIT: An ultra-fast single-node solution for large and complex metagenomics assembly via succinct de Bruijn graph. Bioinformatics 31, 1674–1676. 10.1093/bioinformatics/btv033.

83. Prjibelski, A., Antipov, D., Meleshko, D., Lapidus, A., and Korobeynikov, A. (2020). Using SPAdes De Novo Assembler. Curr. Protoc. Bioinforma. 70, 1–29. 10.1002/cpbi.102.

84. Wick, R.R., Judd, L.M., Gorrie, C.L., and Holt, K.E. (2017). Unicycler: Resolving bacterial genome assemblies from short and long sequencing reads. PLoS Comput. Biol. 13, 1–22. 10.1371/journal.pcbi.1005595.

85. Uritskiy, G. V., DiRuggiero, J., and Taylor, J. (2018). MetaWRAP—a flexible pipeline for genome-resolved metagenomic data analysis. Microbiome 6, 1–13. 10.1186/s40168-018-0541-1.

86. Nurk, S., Meleshko, D., Korobeynikov, A., and Pevzner, P.A. (2017). MetaSPAdes: A new versatile metagenomic assembler. Genome Res. 27, 824–834. 10.1101/gr.213959.116.

87. Institut Pasteur (2024). fq2dna. GitLab. https://gitlab.pasteur.fr/GIPhy/fq2dna.

88. Alneberg, J., Bjarnason, B.S., De Bruijn, I., Schirmer, M., Quick, J., Ijaz, U.Z., Lahti, L., Loman, N.J., Andersson, A.F., and Quince, C. (2014). Binning metagenomic contigs by coverage and composition. Nat. Methods 11, 1144–1146. 10.1038/nmeth.3103.

89. Wu, Y.W., Simmons, B.A., and Singer, S.W. (2016). MaxBin 2.0: An automated binning algorithm to recover genomes from multiple metagenomic datasets. Bioinformatics 32, 605–607. 10.1093/bioinformatics/btv638.

90. Kang, D.D., Li, F., Kirton, E., Thomas, A., Egan, R., An, H., and Wang, Z. (2019). MetaBAT 2: An adaptive binning algorithm for robust and efficient genome reconstruction from metagenome assemblies. PeerJ 2019, 1–13. 10.7717/peerj.7359.

91. Nissen, J.N., Johansen, J., Allesøe, R.L., Sønderby, C.K., Armenteros, J.J.A., Grønbech, C.H., Jensen, L.J., Nielsen, H.B., Petersen, T.N., Winther, O., et al. (2021). Improved metagenome binning and assembly using deep variational autoencoders. Nat. Biotechnol. 39, 555–560. 10.1038/s41587-020-00777-4.

92. Parks, D.H., Imelfort, M., Skennerton, C.T., Hugenholtz, P., and Tyson, G.W. (2015). CheckM: Assessing the quality of microbial genomes recovered from isolates, single cells, and metagenomes. Genome Res. 25, 1043–1055. 10.1101/gr.186072.114.

93. Eren, A.M., Kiefl, E., Shaiber, A., Veseli, I., Miller, S.E., Schechter, M.S., Fink, I., Pan, J.N., Yousef, M., Fogarty, E.C., et al. (2021). Community-led, integrated, reproducible multi-omics with anvi’o. Nat. Microbiol. 6, 3–6. 10.1038/s41564-020-00834-3.

94. Evans, J.T., and Denef, V.J. (2020). To Dereplicate or Not To Dereplicate? mSphere 5, 10–128. 10.1128/msphere.00971-19.

95. Kim, D., Park, S., and Chun, J. (2021). Introducing EzAAI: a pipeline for high throughput calculations of prokaryotic average amino acid identity. J. Microbiol. 59, 476–480. 10.1007/s12275-021-1154-0.

96. Chaumeil, P.A., Mussig, A.J., Hugenholtz, P., and Parks, D.H. (2020). GTDB-Tk: A toolkit to classify genomes with the genome taxonomy database. Bioinformatics 36, 1925–1927. 10.1093/bioinformatics/btz848.

97. Seemann, T. (2014). Prokka: Rapid prokaryotic genome annotation. Bioinformatics 30, 2068–2069. 10.1093/bioinformatics/btu153.

98. Sayers, E.W., Cavanaugh, M., Clark, K., Ostell, J., Pruitt, K.D., and Karsch-Mizrachi, I. (2020). GenBank. Nucleic Acids Res. 48, D84–D86. 10.1093/nar/gkz956.

99. Kodama, Y., Shumway, M., and Leinonen, R. (2012). The sequence read archive: Explosive growth of sequencing data. Nucleic Acids Res. 40, 2011–2013. 10.1093/nar/gkr854.

100. Chen, I.M.A., Chu, K., Palaniappan, K., Pillay, M., Ratner, A., Huang, J., Huntemann, M., Varghese, N., White, J.R., Seshadri, R., et al. (2019). IMG/M v.5.0: An integrated data management and comparative analysis system for microbial genomes and microbiomes. Nucleic Acids Res. 47, D666–D677. 10.1093/nar/gky901.

101. Altschul, S.F., Gish, W., Miller, W., Myers, E.W., and Lipman, D.J. (1990). Basic local alignment search tool. J. Mol. Biol. 215, 403–410. 10.1016/S0022-2836(05)80360-2.

102. Johnson, M., Zaretskaya, I., Raytselis, Y., Merezhuk, Y., McGinnis, S., and Madden, T.L. (2008). NCBI BLAST: a better web interface. Nucleic Acids Res. 36, 5–9. 10.1093/nar/gkn201.

103. SRA Toolkit Development Team (2022). SRA Toolkit. https://trace.ncbi.nlm.nih.gov/Traces/sra/sra.cgi?view=software.

104. Katoh, K., Kuma, K.I., Toh, H., and Miyata, T. (2005). MAFFT version 5: Improvement in accuracy of multiple sequence alignment. Nucleic Acids Res. 33, 511–518. 10.1093/nar/gki198.

105. Waterhouse, A.M., Procter, J.B., Martin, D.M.A., Clamp, M., and Barton, G.J. (2009). Jalview Version 2-A multiple sequence alignment editor and analysis workbench. Bioinformatics 25, 1189–1191. 10.1093/bioinformatics/btp033.

106. Huson, D.H., and Bryant, D. (2006). Application of phylogenetic networks in evolutionary studies. Mol. Biol. Evol. 23, 254–267. 10.1093/molbev/msj030.

107. Shen, W., Le, S., Li, Y., and Hu, F. (2016). SeqKit : A Cross-Platform and Ultrafast Toolkit for FASTA/Q File Manipulation. PLoS One 11, e0163962. 10.1371/journal.pone.0163962.

108. Minh, B.Q., Schmidt, H.A., Chernomor, O., Schrempf, D., Woodhams, M.D., Von Haeseler, A., Lanfear, R., and Teeling, E. (2020). IQ-TREE 2: New Models and Efficient Methods for Phylogenetic Inference in the Genomic Era. Mol. Biol. Evol. 37, 1530–1534. 10.1093/molbev/msaa015.

109. Tria, F.D.K., Landan, G., and Dagan, T. (2017). Phylogenetic rooting using minimal ancestor deviation. Nat. Ecol. Evol. 1, 1–7. 10.1038/s41559-017-0193.

110. Letunic, I., and Bork, P. (2016). Interactive tree of life (iTOL) v3: an online tool for the display and annotation of phylogenetic and other trees. Nucleic Acids Res. 44, W242–W245. 10.1093/nar/gkw290.

111. Suyama, M., Torrents, D., and Bork, P. (2006). PAL2NAL: Robust conversion of protein sequence alignments into the corresponding codon alignments. Nucleic Acids Res. 34, W609–W612. 10.1093/nar/gkl315.

112. Yang, Z. (2007). PAML 4: Phylogenetic analysis by maximum likelihood. Mol. Biol. Evol. 24, 1586–1591. 10.1093/molbev/msm088.

113. Emms, D.M., and Kelly, S. (2019). OrthoFinder: Phylogenetic orthology inference for comparative genomics. Genome Biol. 20, 1–14. 10.1186/s13059-019-1832-y.

114. Harrison, K.J., Crécy-Lagard, V. De, and Zallot, R. (2018). Gene Graphics: A genomic neighborhood data visualization web application. Bioinformatics 34, 1406–1408. 10.1093/bioinformatics/btx793.

