## Supplemental information for "The ancient evolution of far-red light photoacclimation in cyanobacteria"

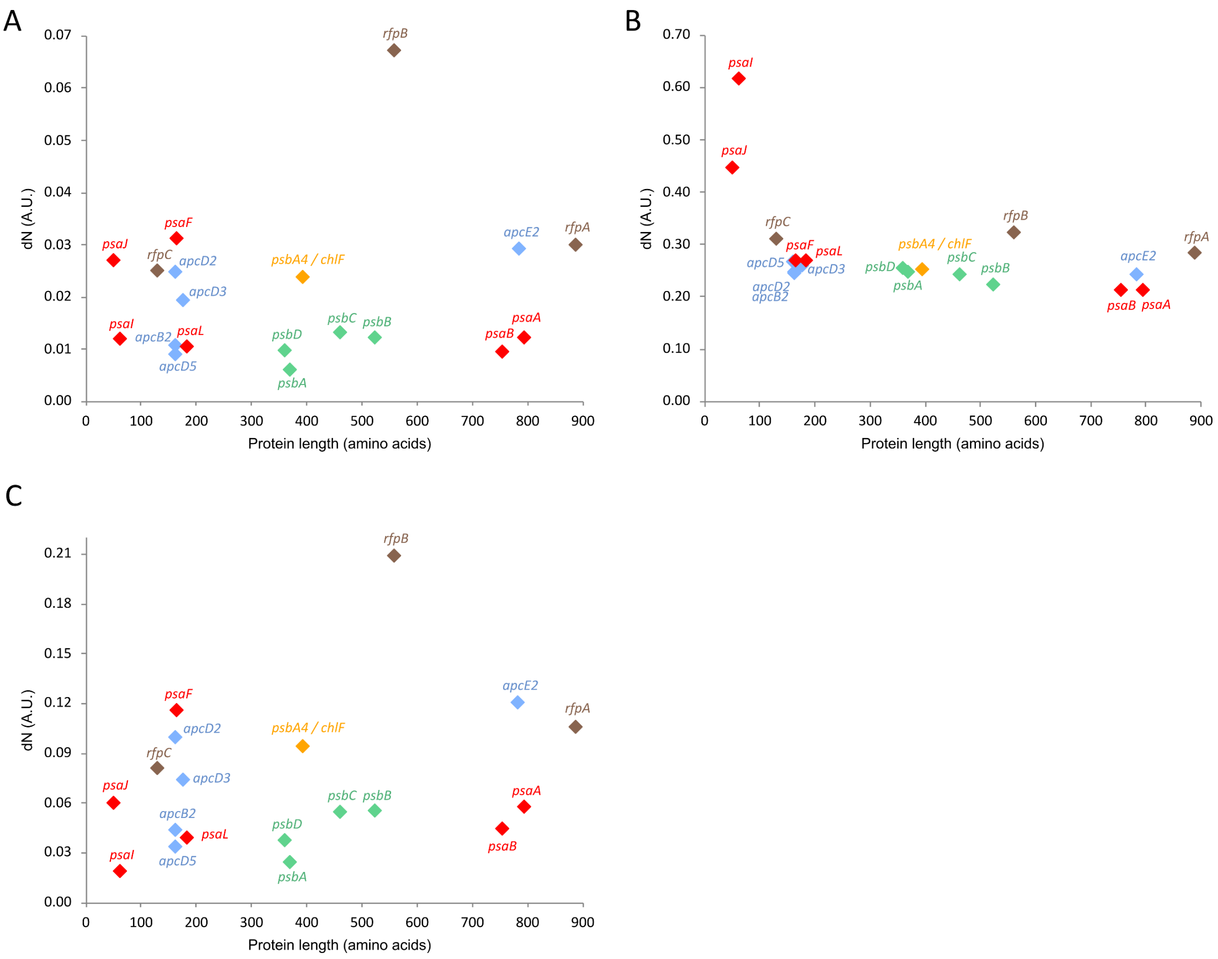


**Figure S1.** Evolutionary rates of FaRLiP core genes. They can be expressed in terms of (A) non-synonymous substitutions (dN), (B) synonymous substitutions (dS) or (C) their ratio (dN/dS). Note the difference in y-axis scales. In general, a higher value on the y-axis means a faster evolutionary rate. Color scheme: PSI genes (red), PSII genes (green), chlorophyll *f* synthase (orange), phycobilisome components (blue) and phytochrome regulatory cascade (brown). Some groups of genes (e.g. PSI) include both fast- and slow-evolving sequences. Dataset based on >80 complete clusters.

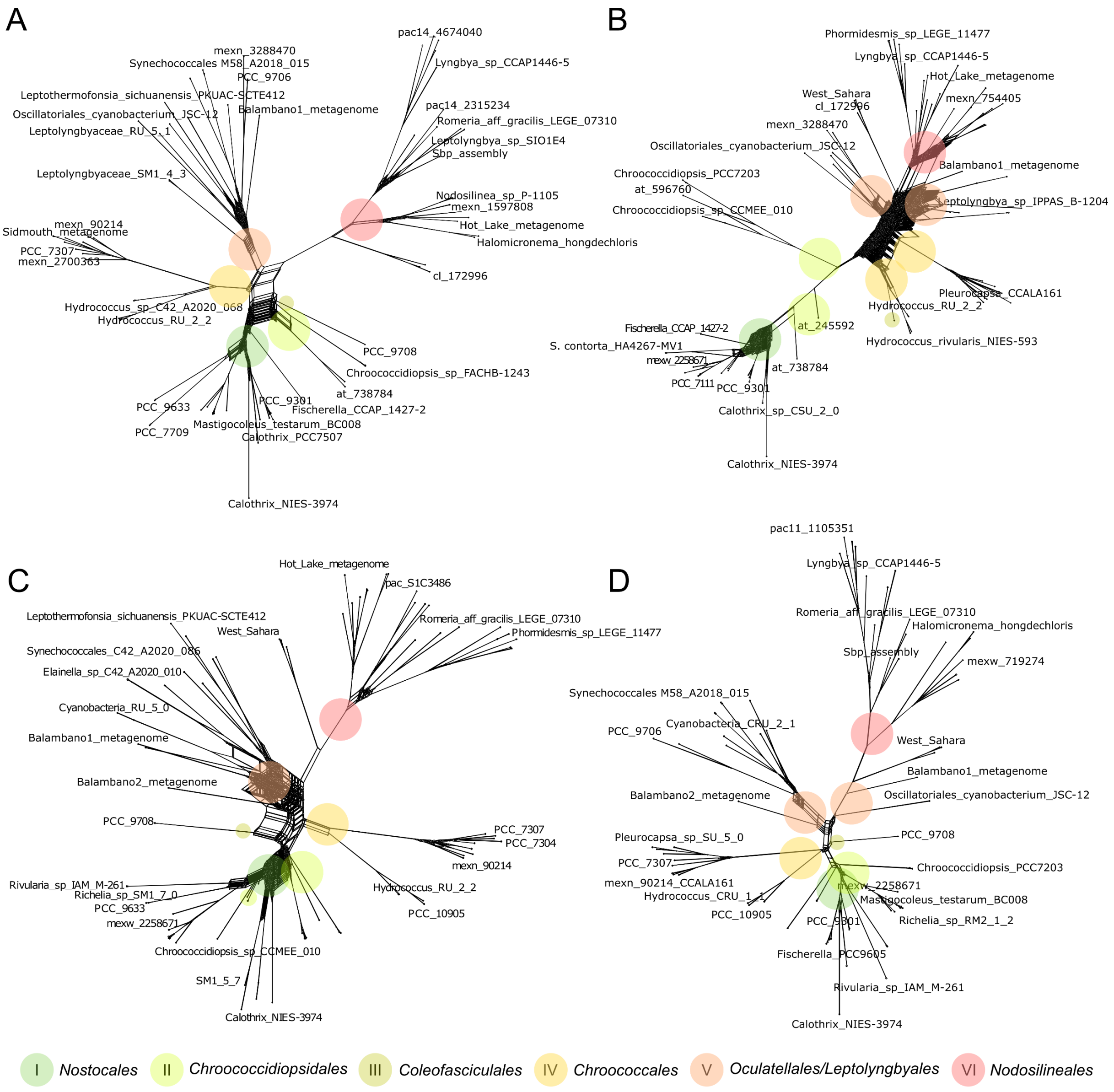


**Figure S2.** Network visualizations of FaRLiP genes. These networks have been constructed by combining multiple DNA trees for: (A) phycobilisome genes *apcE2* and *apcD2/3/5*, (B) Photosystem I paralogues *psaA2/B2/F2/I2/J2/L2*, (C) Photosystem II paralogues *psbA3/B2/C2/D3* – as well as the divergent chlorophyll *f* synthase *chlF* (*psbA4*) – (D) and phytochrome signaling cascade genes *rfpA/B/C*. They approximate how similar gene trees are to each other. Colored circles mark large cyanobacterial groups as shown at the bottom of the figure. Though the PSI gene network is less well resolved, it can be seen that, overall, large splits are very similar between these six gene groups. Approx. 80 taxa used for each gene tree (not all shown). Short metagenome labels are listed in Table S12. Protein tree networks were also constructed, and were very similar to those shown above.

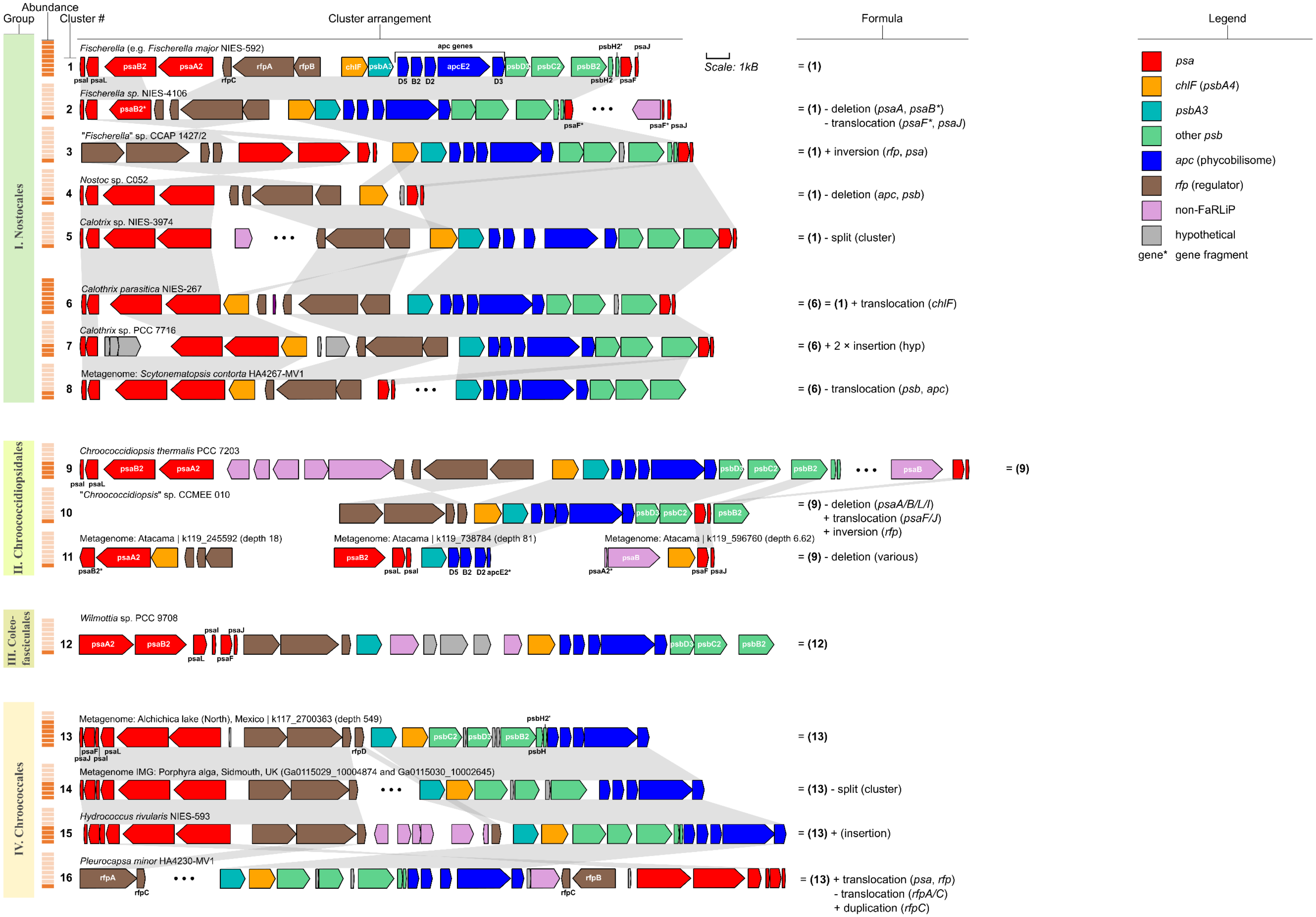


**Figure S3.** *Diversity of FaRLiP cluster gene arrangements within the cyanobacteria. (Figure continues below)*

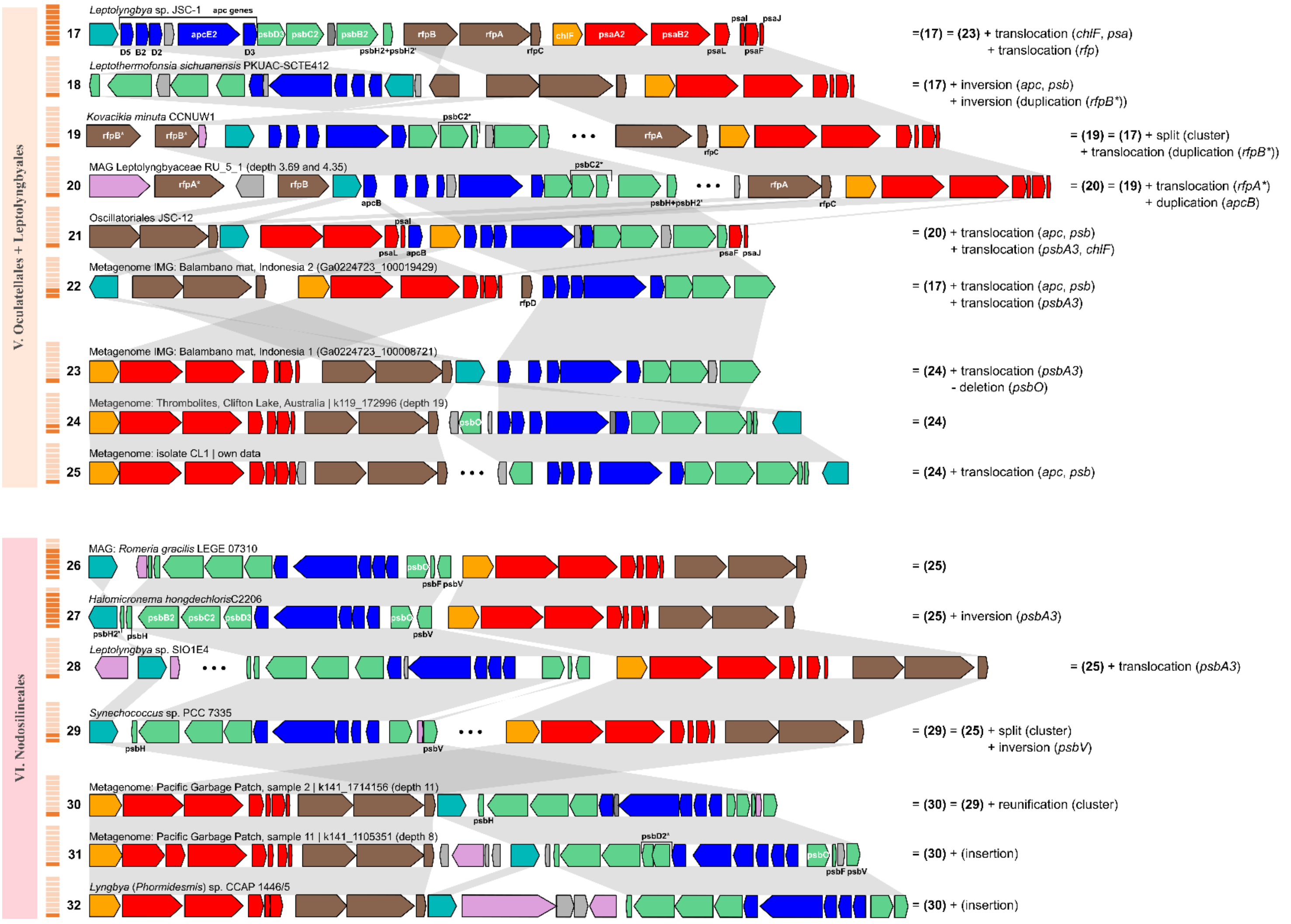


**Figure S3.** *Diversity of FaRLiP cluster gene arrangements within the cyanobacteria. (Figure legend continues below)*

**Figure S3.** Diversity of FaRLiP cluster gene arrangements within the cyanobacteria *(Figure continues above)*. FaRLiP is present within six cyanobacterial groups (left): *Nostocales* (I), *Chroococcidiopsidales* (II), *Coleofasciculales* (III), *Chroococcales* (IV), *Oculatellales* + *Leptolyngbyales* (V – grouped together due to polyphyly in the gene tree) and *Nodosilineales* (VI). Every cluster shape was labeled with a number (from 1 to 32), shown under the column 'Cluster #'. The 'Abundance' column lists how many times a cluster shape occurs. Each orange bar indicates the presence of the respective shape in one strain or metagenome, capped at a maximum of 8. Grey highlights between clusters mark homologous regions. Within groups, rare clusters typically represent variants of more abundant clusters (clusters 2-5 as variants of cluster 1). Formulas on the right-hand side suggest necessary mutations. However, in some lineages (*Chroococcales* – IV), multiple variants may have similar abundances, and it remains unclear which was the ancestral one (clusters 13-15). Lineage V (*Oculatellales* + *Leptolyngbyales*) shows the highest syntenic diversity, with multiple mutations necessary to explain variants. There are indications that the most common arrangement (17) may not have been the ancestral one. Instead, clusters 23-25 may better reflect the ancestor of group V due to the maintenance of traits characterizing the earlier-branching group VI (*Nodosilineales*): the order of *chlF*, then *psa*, then *rfp*, as well as the presence of *psbO*.

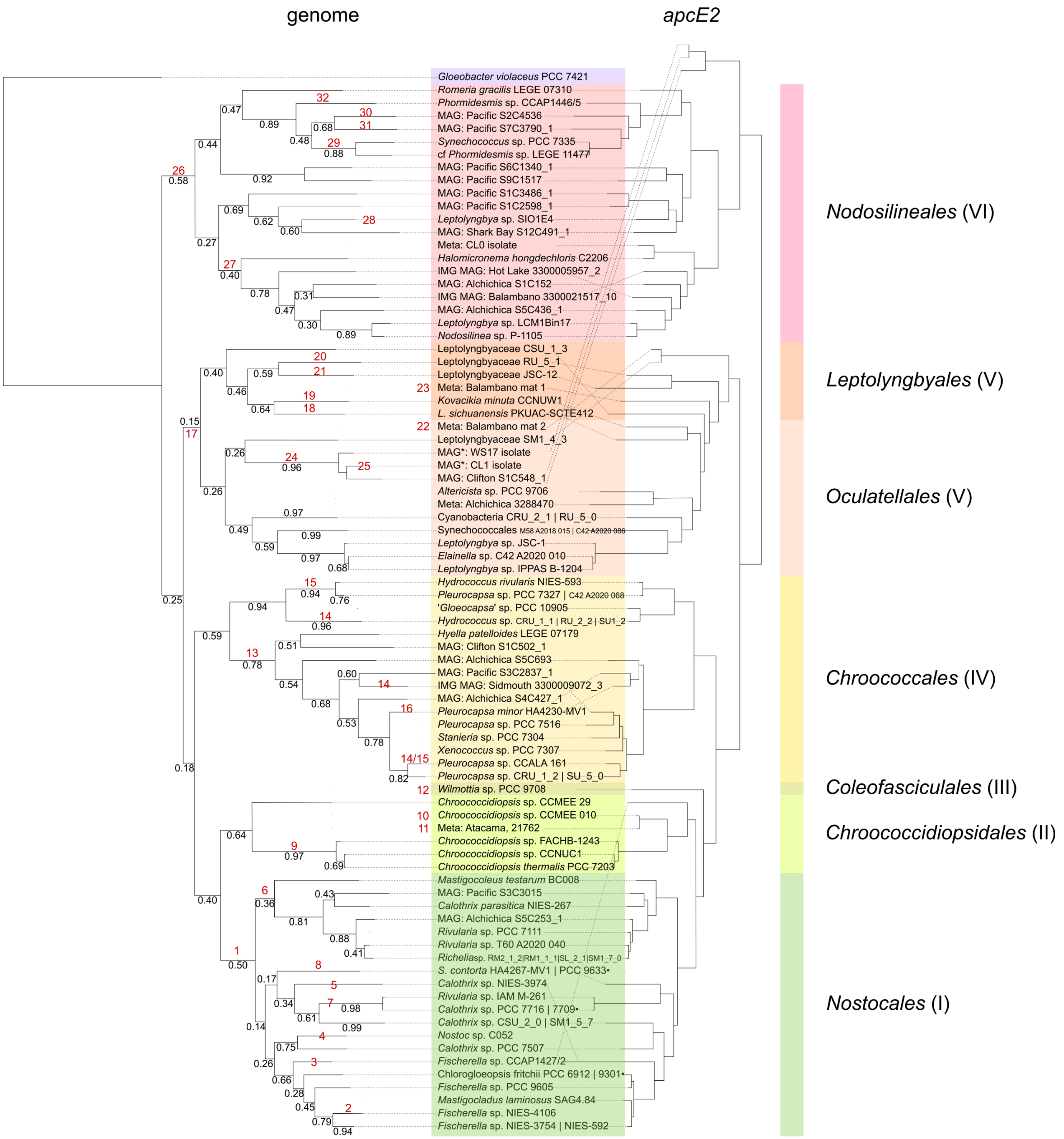


**Figure S4.** Gene arrangement within the FaRLiP cluster is conserved in accordance with the phylogenetic tree. The evolutionary tree of FaRLiP cyanobacteria is split into six groups. These are color-coded and marked on the right-hand side (I – VI). A genome tree is available for most of the strains (left); where a genome is not available, strain relationships can be approximated from the *apcE2* phylogeny (right). Gene arrangements are listed as red numbers, and are shown in detail in Figure S3. Mutations between cluster shapes listed in Figure S3 can be observed to be vertically inherited in the tree. Where known, particular gene arrangements are listed above the most recent common ancestor to include them. Different cluster shapes can be present in only one known lineage (e.g. cluster 3) or more (e.g. cluster 6). Some clusters (12, 22, 23) do not have an associated genome. Two genomes (*Chroococcdiopsis* sp. CCMEE 29 and the root *Gloeobacter violaceus* PCC 7421) were added to clarify the phylogeny, and do not have an associated FaRLiP cluster.

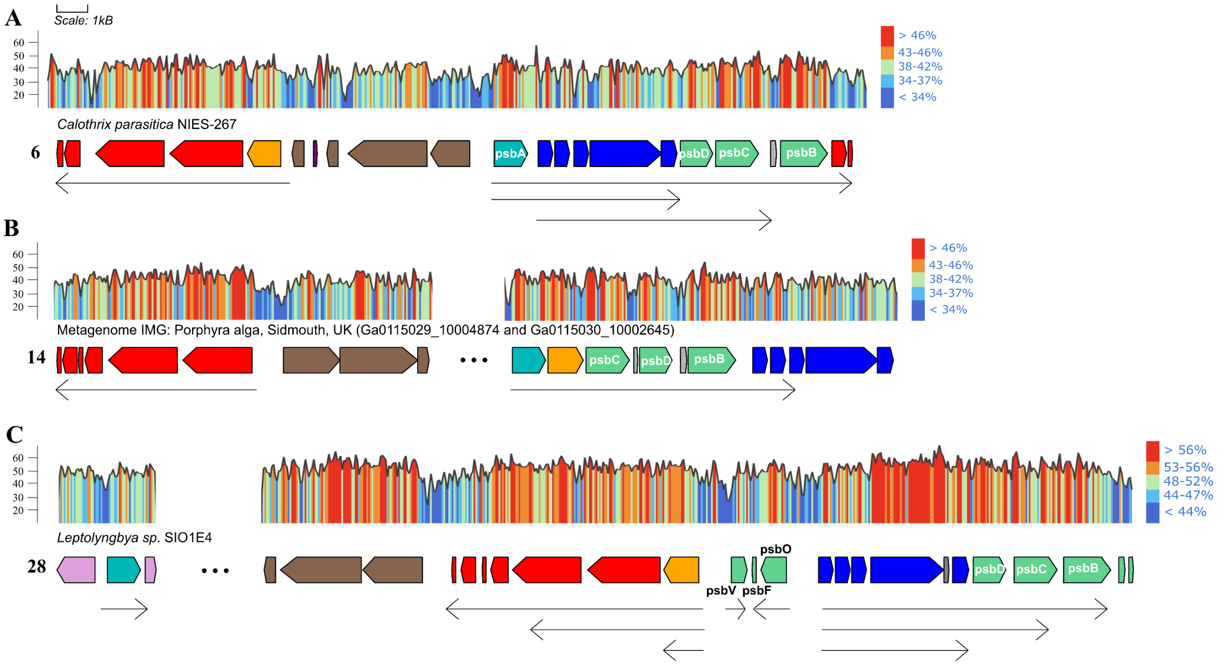


**Figure S5.** Metatranscriptomics data of the FaRLiP cluster. Reads from transcriptomic datasets were assembled in previous studies to recover mRNAs. These may include multiple genes, and are equivalent to putative operons (shown as arrows below the clusters). The examples above were recovered from metatranscriptomic assemblies for strains closely related to those containing cluster types 6 (A), 14 (B) and 28 (C). See Figure S3 for all cluster types; see Table S10 for data sourcing. Naming is based on gene homology and cluster structure. GC% of the gene clusters is shown above them (red high, blue low, according to the color schemes on the right). Note, intergenic regions are AT-rich. Gene color scheme: *psa* (red); *psb* (green), except for *psbA3* (turquoise); *chlF* (yellow); *apc* (blue); *rfp* (brown).

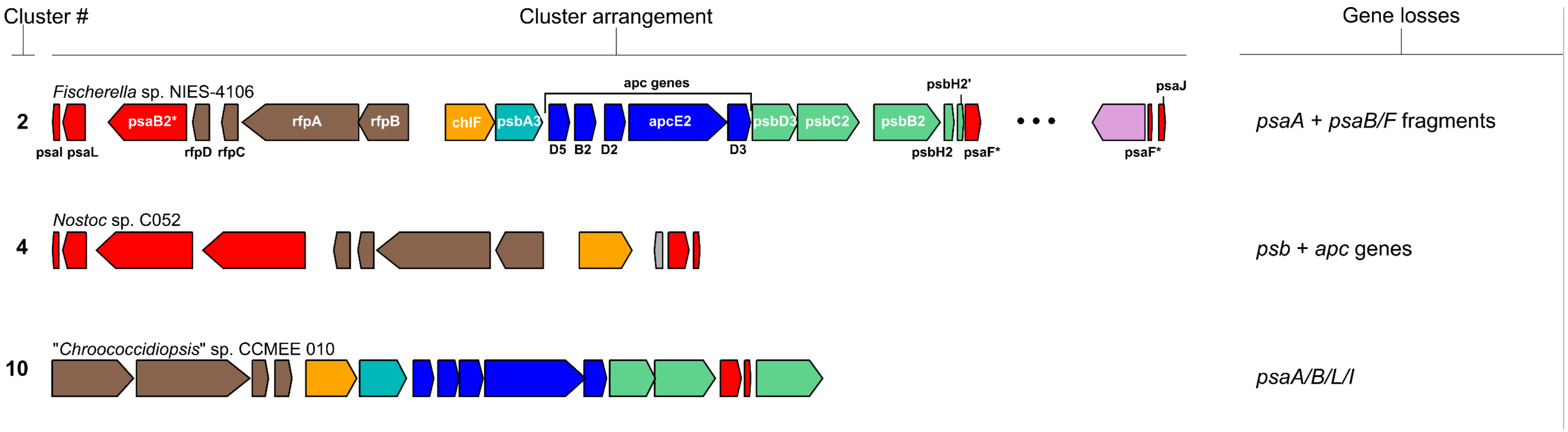


**Figure S6.** Partial FaRLiP clusters. As a result of gene loss, these clusters contain fewer than the typical 19 core genes. One of them, from '*Chroococcidiopsis*' sp. CCME 010, was proven to be functional[1]. There appears to be a tendency for genes in the same operon, or with related functional roles, to be lost together, *psa* genes for clusters 2 and 10, *psb* and *apc* genes for cluster 4. The numbers correlate with Figure S3.

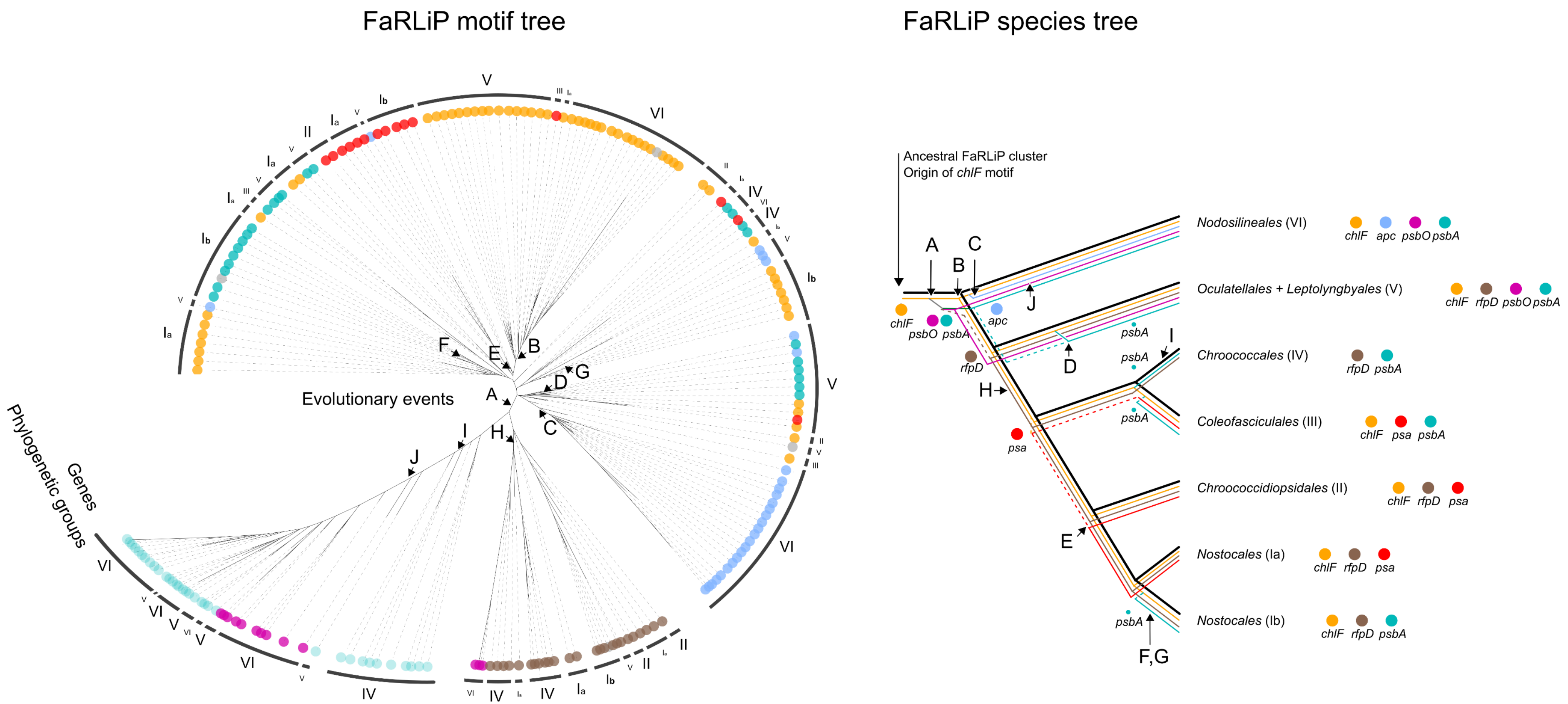


**Figure S7.** FaRLiP motif evolution supports vertical descent. *(Figure continues below)*

### Evolutionary events

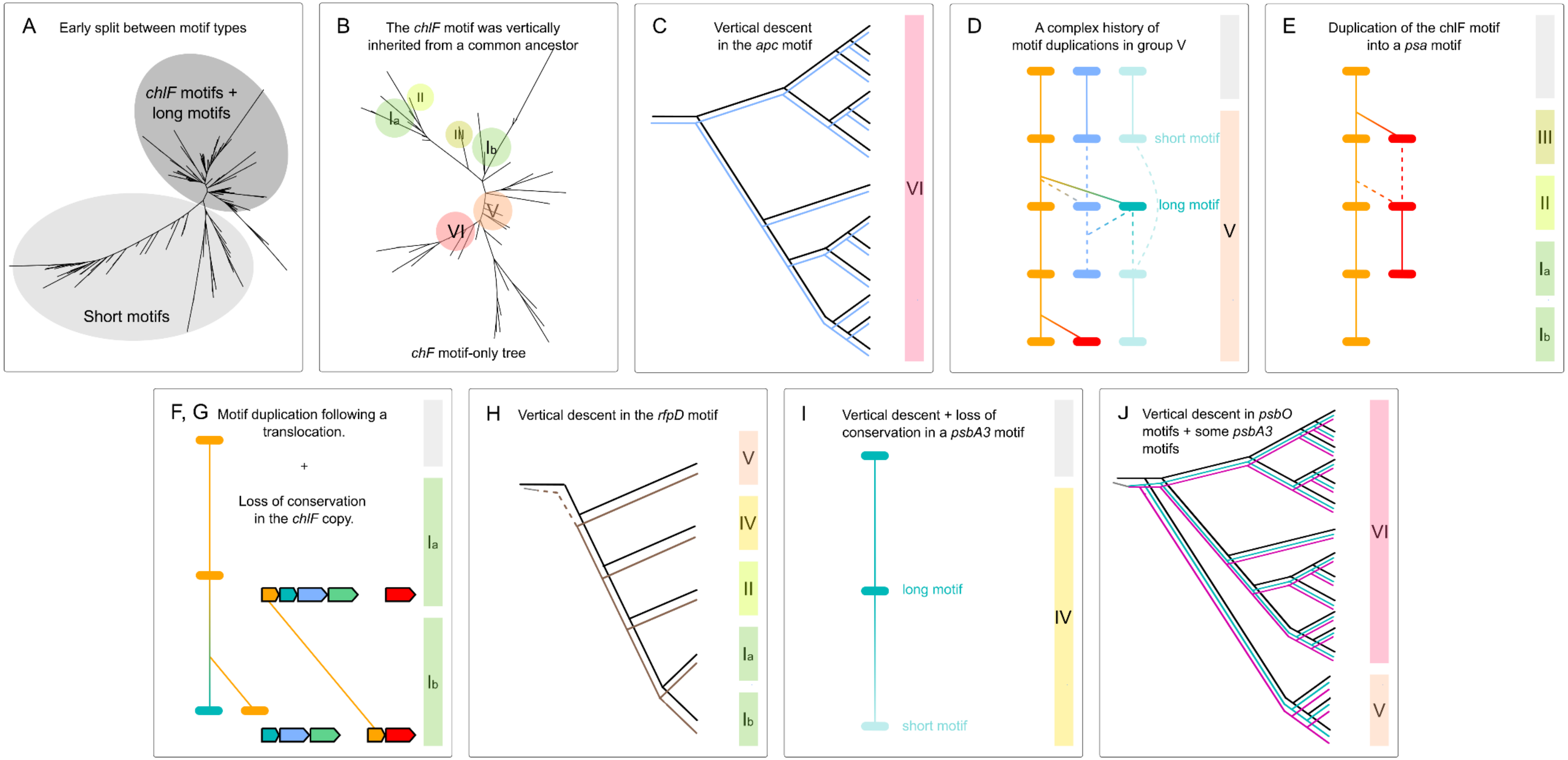


**Figure S7.** FaRLiP motif evolution supports vertical descent. The evolution of the FaRLiP motif is complex, with numerous duplications leading to orthologs and paralogues. Interpretations need to not only consider the motif tree (top left) but also the species tree (top right) as well as motif conservation (Figure 5A). However, certain evolutionary events are clear. (A) Before the FaRLiP cluster MRCA, a duplication occurred, leading to the ancestor of many short motifs on one side, and all long/*chlF* motifs on the other. This split is visible, though much noisier, when only the 33 bp of the conserved 3' region are considered. The ancestral sequence was likely a *chlF* motif, due to its high conservation. (B) The *chlF* motif was inherited vertically. Motifs from the same phylogenetic groups cluster together, in a way that matches the species tree. The mismatch of group $I_b$ will be discussed later. (C) The motif in front of *apc* genes was inherited vertically within group VI. (D) Group V is characterized by numerous duplications leading to motif paralogues. Sometimes, these homologous relationships are clear (solid lines), but most often they are uncertain (dashed lines). Supporting evidence can include strong conservation (for the duplication of a *chlF* motif into a *psa* or *psbA3* motif) or cluster arrangement (for the maintenance of an *apc* and short *psbA3* motif in samples WS17/CL1). (E) Vertical descent is observed within *psa* motifs in groups $I_a$ and II, after this motif duplicated from the *chlF* motif in their common ancestor. It is uncertain whether the *psa* motif of group III represents an independent occurrence. (F) In group $I_b$, the highly conserved *chlF* motif was maintained in front of the *psbA3* gene after a translocation of the *chlF* gene. (G) In contrast, the motif translocated together with the *chlF* gene appears to have lost its highly conserved nature, explaining its divergent position in the phylogeny. (H) The motif in front of *rfpD* was likely inherited vertically. (I) Short *psbA3* motifs in group IV appear to represent recent losses from long *psbA3* motifs. Despite the motif phylogeny suggesting a close relationship to more ancient short motifs instead, this is not supported by cluster structure, and it highlights the need to consider multiple sources of information when working with short sequences. (J) Evidence supports vertical inheritance of short *psbA3* motifs and *psbO* motifs within group VI and part of group V.

| Name | Culture collection[a] | Equivalent | Medium[b] | Category[c] | Sampling site |
|---|---|---|---|---|---|
| *Altericista* sp. | PCC 9706 | | $BG11_0$ + $NaNO_3$ (2 mM) + $NaHCO_3$ (10 mM) (S) | Other | River Seine, Paris area, France |
| *Calothrix* sp. | PCC 7709 | CALU 653, CCALA 033, CCAP 1410/6, DCC D0253a | $BG11_0$ | Other | Tree bark, near river Rio Arimao, Las Villas, Cuba |
| *Calothrix* sp. | PCC 9633 | | $BG11_0$ + $NaHCO_3$ (5 mM) (S) | Other | Retention basin of river Oise (Paris area), France |
| *Chlorogloeopsis fritschii* | PCC 9301 | | $BG11_0$ + $NaHCO_3$ (5 mM) (S) | Unknown | Possibly Spain |
| *Chroococcidiopsis* sp. | PCC 7432 | CCALA 042 | $BG11_0$ + $NaHCO_3$ (5 mM) (S) | Other | Spring near Santa Fe, Cuba |
| *Chroococcidiopsis* sp. | PCC 7434 | CCALA 041 | $BG11_0$ + $NaHCO_3$ (5 mM) (S) | Other | Pool, botanical garden, Havana, Cuba |
| *Chroococcidiopsis* sp. | PCC 7439 | CCALA 046 | $BG11_0$ + $NaHCO_3$ (5 mM) (S) | Other | Sand beach near Mamaia, Constanța, Romania |
| *Fischerella* sp. | CCAP 1427/2 | | JM | Other | Plant material, Cuba |
| *Gloeocapsa* sp. | PCC 10905 | | BG11 + $NaHCO_3$ (5 mM) (S) | Unknown | Hot spring, south of France |
| *Mastigocladus* sp. | SAG 4.84 | | BG11 | Other | Thermal spring of Reyhjanes/Isafjord, Iceland |
| *Phormidesmis* sp. | CCAP 1446/5 | | ASW:BG | Other | Sea off Lowestoft, Suffolk, England, UK |
| *Pleurocapsa* sp. | PCC 7516 | | ASNIII + B12 (10 µg $L^{-1}$) | Other | Rock chip, station B, I'Ile Riou, Marseille, France |
| *Rivularia* sp. | PCC 7111 | | $BG11_0$ (95%) + 4X Sea Water solution (5%) | Microbial mat | Hemispherical macroscopic colony, intertidal, Bodega, California, USA |
| *Stanieria* sp. | PCC 7304 | | ASNIII + $NaHCO_3$ (5 mM) + B12 (10 µg $L^{-1}$) (S) | Other | Epiphyte on *Rhodochorton* sp., high intertidal zone, Bodega Marine Laboratory, California, USA |
| *Wilmottia* sp. | PCC 9708 | | $BG11_0$ + $NaNO_3$ (2 mM) + $NaHCO_3$ (10 mM) (S) | Other | Retention basin of river Oise (Paris area), France |
| *Xenococcus* sp. | PCC 7307 | ATCC 29375 | ASNIII + $NaHCO_3$ (5 mM) + B12 (10 µg $L^{-1}$) (S) | Other | Rock chip, high intertidal zone, Horseshoe Cove, Bodega Marine Laboratory, California, USA |

**Table S2**. Strains available from culture collections whose genetic data was used in this study. [a] Culture collections: CCAP / SAMS (Culture Collection of Algae and Protozoa at the Scottish Association for Marine Science, Oban, Scotland), SAG (Culture Collection of Algae at Göttingen University, Göttingen, Germany), PCC (Pasteur Culture of Cyanobacteria, Paris, France). The same strains may be/have been present in different culture collections, with alternative codes (see 'Equivalent' column). [b] JM – Jaworski's Medium[2], ASW:BG – ASWIII and BG11 medium for marine cyanobacteria[3], BG11 – standard cyanobacterial growth medium[4]. $BG11_0$ – BG11 without sodium nitrate. (S) marks solid media (0.9% w/v agar). [c] Environments were categorized as per Figure 1.

| Sampling site | Category | Label | Order | Completion (%) | Redundancy (%) | NCBI MAG Accession | SRA Accession | Data source |
|---|---|---|---|---|---|---|---|---|
| Alchichica lake stromatolites, Mexico | Microbialite | Alchichica S1C152 | *Nodosilineales* | 97.87 | 1.63 | | SRR3311162 | [5] |
| | | Alchichica S5C436_1 | *Nodosilineales* | 97.83 | 2.59 | | SRR3311151/3/6 | |
| | | Alchichica S5C693 | *Chroococcales* | 96 | 0.66 | | SRR3311151/3/6 | |
| | | Alchichica S4C427_1 | *Chroococcales* | 96.58 | 0.66 | | SRR3310977 | |
| | | Alchichica S5C253_1 | *Nostocales* | 82.5 | 4.6 | | SRR3311151/3/6 | |
| Sebkha Oum Dba, Morocco | Microbial mat | WS17 isolate | *Oculatellales* | 98.55 | 2.45 | | N/A | Own data |
| Clifton lake thrombolites, Australia | Microbialite | CL1 isolate | *Oculatellales* | 94.75 | 1.36 | | N/A | |
| | | Clifton S1C502_1 | *Chroococcales* | 83.84 | 0.66 | | SRR3280822 | [6] |
| Pacific Garbage Patch microplastics | Microbial mat | Pacific S2C4536 | *Nodosilineales* | 94.7 | 3.22 | | SRR3401470 | [7] |
| | | Pacific S7C3790_1 | *Nodosilineales* | 97.01 | 1.9 | | SRR3401475 | |
| | | Pacific S6C1340_1 | *Nodosilineales* | 99.46 | 1.9 | | SRR3401474 | |
| | | Pacific S9C1517 | *Nodosilineales* | 96.6 | 1.9 | | SRR3401477 | |
| | | Pacific S1C3486_1 | *Nodosilineales* | 97.46 | 1.09 | | SRR3401469 | |
| | | Pacific S1C2598_1 | *Nodosilineales* | 97.46 | 3.26 | | SRR3401469 | |
| | | Pacific S3C2837_1 | *Chroococcales* | 88.81 | 3.54 | | SRR3401471 | |
| | | Pacific S3C3015 | *Nostocales* | 83.35 | 0.7 | | SRR3401471 | |
| Shark Bay stromatolites, Australia | Microbialite | Shark Bay S12C491_1 | *Nodosilineales* | 69.52 | 2.36 | | SRR6462995 | [8] |

**Table S3.** Metagenome-assembled genomes (MAGs) built and used in this study. Completion and redundancy values were calculated with CheckM following bin refinement by anvi'o. Taxonomic assignment was executed with GTDB-Tk and cross-checked with recent literature.

| Code | IMG scaffold code | Category | Sampling location | Notes |
|---|---|---|---|---|
| Atacama_gypsum[a] | Ga0395813_0004 | Other | Gypsum rocks, Atacama Desert, Chile | [9] |
| Balambano1_metagenome | Ga0224723_1000087 | Microbial mat | Balambano mat, Indonesia | Permission kindly granted by Rachel Simister and Sean Crowe (University of British Columbia, Canada). Similar sample published[10]. |
| Balambano2_metagenome | Ga0224723_1000194 | Microbial mat | Balambano mat, Indonesia | Permission kindly granted by Rachel Simister and Sean Crowe (University of British Columbia, Canada). Similar sample published[10]. |
| Hot_Lake_metagenome | Ga0081520_100035 | Microbial mat | Biofilm, Hot Lake, Washington, USA | Permission kindly granted by Stephen Lindermann (Purdue University, USA). |
| Sidmouth_metagenome | Ga0115030_1000264, Ga0115029_1000487 | Other | *Porphyra* algae metagenome, Sidmouth, UK | Permission kindly granted by Susan Brawley (University of California, USA). Dataset produced as part of the *Porphyra umbilicalis* genome project[11] under US Department of Energy Contract DE-AC02-05CH11231. |

**Table S4.** A subset of FaRLiP clusters in this study were recovered from the IMG database. Only complete clusters (or, in the case of the Sidmouth sequences, complete naturally split clusters) were selected from the larger dataset previously discussed in Antonaru et al., 2020. For the complete dataset, please see Tables S3 and S4 in the aforementioned work. [a] Only used in the environmental analysis due to high similarity with other sequences.

| Protein | Consensus | Position | Role |
|---|---|---|---|
| **ApcE2** | YAIVAGDGSILSANVRGLRGVIPEDVTEATIVALRAMRRQSLDYFL | 184 – 229 | FR-specific, non-covalent binding of bilin chromophore[13] |
| **PsbA4** | YQDREWELSYRLGMRPWISLAFT | 148 – 170 | QD site is necessary and sufficient for chl *f* synthesis[14] |
| **PsbA3** | AFHYIPALCCYLGREW | 117 – 132 | Y120 has been hypothesized to be associated with the binding of chlorophyll *d* as $Chl_{D1}$ [15–17]. Nearby residues are likely to stabilize this interaction. |
| **PsbB2** | MHNALCAGFAGSML | 25 – 38 | Part of the first transmembrane helix of PsbB2. H26 is a chlorophyll-binding residue. F33 is hypothesized to be involved in the binding of chl *f* (F34 in citation)[15]. |
| **PsbC2** | YHSLKGPEKL_AGFFBFDWSDKDKVTQILG | 127 – 155 | A FaRLiP-specific deletion is in the stromal loop between transmembrane helices 2 and 3. Based on genetic data, it has been suggested to influence allophycocyanin binding[17]. |
| **PsbH2** | QPKKVAPVQYLLRNFNSEAGKVT | 7 – 29 | Mutations seem to affect the stromal-side helix and loop. Based on genetic data, the FR-conserved AP and QY may coordinate chl *a* 617[17]. |
| **PsaA2** | SHLAWVCQFLGFHSFAMY | 445 – 462 | Unknown.<br>Represents the majority of transmembrane helix 7. |
| **PsaB2** | TPLS FGYWKDKPVALSIV | 692 – 709 | Unknown.<br>Has been suggested as a chlorophyll *f* binding site based on electrostatic potential[18]. |
| **PsaI2** | PWIMIPLVFYILPFPIFA | 17 – 34 | The conserved Y25 has been suggested as a chl *f* binding site based on electrostatic potential[18]. |

**Table S5.** Conserved motifs in proteins involved in far-red photoacclimation. FaRLiP-specific residues are underlined. Sequences may not fully represent a classical consensus, and occasionally incorporate rare amino acid variants in order to recover as many FaRLiP sequences as possible. Position within the protein is given for *Chroococcidiopsis thermalis* PCC 7203.

| Function | Identifier | iTOL code (gene) | iTOL code (protein) |
|---|---|---|---|
| Phycobilisome | *apcB2* | 1301336668256591707756200 | 130133666831741711377234 |
| | *apcD2* | 1301336668256611707756200 | 130133666831771711377234 |
| | *apcD3* | 1301336668256631707756200 | 130133666831791711377235 |
| | *apcD5* | 1301336668248981707756155 | 130133666831701711377234 |
| | *apcE2* | 1301336668256661707756201 | 130133666831811711377235 |
| Photosystem I | *psaA2* | 1301336668256691707756201 | 130133666831841711377235 |
| | *psaB2* | 1301336668256721707756201 | 130133666831861711377235 |
| | *psaF2* | 1301336668256741707756201 | 130133666831881711377236 |
| | *psaI2* | 1301336668256761707756201 | 130133666831911711377236 |
| | *psaJ2* | 1301336668256781707756202 | 130133666831931711377236 |
| | *psaL2* | 1301336668256801707756202 | 130133666831981711377236 |
| Chlorophyll *f* synthase | *chlF* (*psbA4*) | 1301336668256941707756203 | 130133666832131711377238 |
| Photosystem II | *psbA3* | 1301336668256821707756202 | 130133666832041711377237 |
| | *psbB2* | 1301336668256841707756202 | 130133666832061711377237 |
| | *psbC2* | 1301336668256871707756203 | 130133666832081711377237 |
| | *psbD3* | 1301336668256901707756203 | 130133666832111711377238 |
| Phytochrome signaling cascade | *rfpA* | 1301336668256961707756203 | 130133666832151711377238 |
| | *rfpB* | 1301336668256981707756203 | 130133666832171711377238 |
| | *rfpC* | 1301336668257001707756204 | 130133666832221711377239 |

**Table S6.** Phylogenetic trees of core FaRLiP genes are available on the interactive Tree of Life (iTOL) webserver, and can be accessed through the associated hyperlinks. Gene trees (left) are very similar to protein trees (right). The format of the hyperlinks is ‘https://itol.embl.de/tree/’iTOL_code. *psbA4* is equivalent to *chlF*.

| Function | Gene | dN | dS | dN / dS | Protein size (aa) |
|---|---|---|---|---|---|
| Phycobilisome | *apcB2* | 0.011 | 0.246 | 0.044 | 161 |
| | *apcD2* | 0.025 | 0.248 | 0.100 | 159 - 182 |
| | *apcD3* | 0.019 | 0.260 | 0.074 | 158 - 214 |
| | *apcD5* | 0.009 | 0.266 | 0.034 | 158 |
| | *apcE2* | 0.029 | 0.242 | 0.121 | 733 - 806 |
| Photosystem I | *psaA2* | 0.012 | 0.212 | 0.058 | 750 - 790 |
| | *psaB2* | 0.009 | 0.213 | 0.044 | 740 - 744 |
| | *psaF2* | 0.031 | 0.269 | 0.116 | 159 - 177 |
| | *psaI2* | 0.012 | 0.617 | 0.019 | 28 - 116 |
| | *psaJ2* | 0.027 | 0.446 | 0.060 | 33 - 82 |
| | *psaL2* | 0.011 | 0.268 | 0.039 | 131 - 204 |
| Chlorophyll *f* synthase | *chlF* (*psbA4*) | 0.006 | 0.247 | 0.024 | 366 - 424 |
| Photosystem II | *psbA3* | 0.024 | 0.251 | 0.095 | 303 - 369 |
| | *psbB2* | 0.012 | 0.222 | 0.055 | 509 - 554 |
| | *psbC2* | 0.013 | 0.242 | 0.055 | 404 - 495 |
| | *psbD3* | 0.010 | 0.254 | 0.038 | 351 - 354 |
| Phytochrome signaling cascade | *rfpA* | 0.030 | 0.283 | 0.106 | 684 - 977 |
| | *rfpB* | 0.067 | 0.322 | 0.209 | 361 - 694 |
| | *rfpC* | 0.025 | 0.310 | 0.081 | 122 - 172 |

**Table S7.** Metrics of evolutionary change in core FaRLiP paralogues. dN – ratio of non-synonymous substitutions; dS – ratio of synonymous substitutions. Although all of the genes described appear to be under purifying selection (dN/dS < 1), some show more change than others. For example, *rfpB*, a component of the signaling cascade, is especially fast-evolving (high dN and dN/dS). Estimates for small proteins are less reliable. Dataset based on >80 complete FaRLiP clusters.

| Label | Putative role | Location | Occurrence (% clusters) | Notes | Approx. size (aa) |
|---|---|---|---|---|---|
| ***rfpD*** | Response regulator. Potentially involved in the phytochrome signaling cascade. REC superfamily. Shown to be preferentially transcribed under FRL[19]. DNA-binding based on DRNApred[20] and NCBI Conserved Domains [21]. Conserved, putative functional residues predicted include N3-R4 (DNA-binding), D58 (phosphorylation site) and V121-A126 (dimer interface), as per *Chroococcidiopsis thermalis* PCC 7203. | Downstream of *rfpC*, same orientation. Occasionally separated by a non-FaRLiP insertion | 35 | Absent in early-branching *Nodosilineales* (VI). | 80 |
| ***psbH2*** | Assembly and/or stability of PSII.[22] | Downstream of *psbB2*, same orientation. | 84 | Absent in the single sequence of group III. | 75 |
| ***psbH2'*** | Unknown. May assist the function of PsbH2. No significant similarity to proteins with a known function. | Downstream of *psbH2*, same orientation. | 71 | 86% of the clusters which contain *psbH2*. Never present without *psbH2*. In group V, *psbH2* and *psbH2'* are connected in a single protein. | 48 |
| ***psbO2*** | Manganese-stabilizing protein[23,24] | Upstream of the FaRLiP allophycocyanin genes, reverse orientation. | 28 | All *Nodosilineales* (VI), 16% of *Oculatellales/ Leptolynbyales* (V) | 275 |
| ***psbV2*** | Manganese-stabilizing protein[23,24] | Either: upstream of *chlF* / *psbA4*, reverse orientation, opposite orientation to *psbF2* and *psbO2*; or (when at the edge of the FaRLiP cluster) downstream of *psbF2*, same orientation | 24 | 87% of clusters including *psbO2* (20/23). | 185 |
| ***psbF2*** | PSII component[24,25] | Between *psbO2* and *psbV2* (downstream of both). Same orientation as *psbO2*. | 17 | 70% of clusters including *psbV* (14/20). | 45 |

**Table S8.** Accessory FaRLiP genes. These genes are found in a subset of far-red clusters, are distinct from standard (white-light) paralogues, and in all cases except for *psbF2* (which lacks data), there is evidence that they are preferentially transcribed or translated under far-red light[19,26–28].

| Species name | Cluster GC content [%] | Genome / contig GC content [%] | Contig size (where applicable) [kbps] |
|---|---|---|---|
| *Fischerella major* NIES-592 | 42.27 | 42.28 | 53 |
| *Fischerella* sp. NIES-3754 | 42.31 | 40.99 | — |
| *Chlorogloeopsis fritchii* PCC 9212 | 43.16 | 41.44 | 233 |
| *Fischerella* sp. NIES-4106 (plasmid) | 41.26 | 41.32 | 300 |
| *Calothrix* sp. PCC 7507 | 44.07 | 42.25 | — |
| *Calothrix* sp. NIES-3974 | 41.27 | 41.53 | — |
| *Calothrix parasitica* NIES-267 | 39.87 | 36.77 | — |
| *Mastigocoleus testarum* BC008 | 40.82 | 38.41 | 77.5 |
| *Chroococcidiopsis thermalis* PCC 7203 | 45.51 | 44.44 | — |
| *Hydrococcus rivularis* NIES-593 | 47.32 | 45.76 | 102 |
| *Pleurocapsa minor* PCC 7327 | 47.52 | 45.19 | — |
| *Leptolyngbya* sp. JSC-1 | 50.54 | 49.12 | 1,530 |
| *Synechococcus* sp. PCC 7335 | 49.60 \| 48.94 [a] | 48.15 | 4,720 |
| *Halomicronema hongdechloris* C2206 | 49.60 | 54.62 | — |

**Table S9.** GC content. With the exception of *Halomicronema hongdechloris* C2206, most FaRLiP clusters have GC values that closely match (±2-3%) that of their host genome. This supports the hypothesis of vertical descent. Where a full genome was not available (marked by —), the contig containing the cluster was used for comparison, and the contig size is listed. [a]*Synechococcus* sp. PCC 7335 has a split FaRLiP cluster. GC percentages for both halves are listed.

| Related species | Group | Cluster shape | Location | Location details | Scaffolds | Permissions and references |
|---|---|---|---|---|---|---|
| *Calothrix parasitica* NIES-267 | I | 6 | Estuarine microbial mat communities from Elkhorn Slough, Moss Landing, CA, USA | | JGI12140J13697_1000045, JGI12140J13697_1000181 | Sample generation and sequencing for this dataset was supported by the LLNL Biofuels Scientific Focus Area SCW1039, and Join Genome Institute Community Sequencing Program award #701 to Jennifer Pett-Ridge (Lawrence Livermore National Laboratory). |
| | | | Microbial communities of thrombolites from Highborne Cay, Bahamas | 0-3 mm depth | Ga0103784_1000008, Ga0103784_1000012, Ga0103785_100006, Ga0103785_100010, Ga0103785_100020 | [29] |
| | | | | 3-5 mm depth | Ga0103786_1000001, Ga0103786_1000004, Ga0103787_1000001, Ga0103787_1000004 | [29] |
| | | | | 5-9 mm depth | Ga0103788_10000045, Ga0103788_1000010, Ga0103788_1000023 | [29] |
| *Chroococcales* cyanobacterium | IV | 14 | *Siderastrea* coral microbial communities from Puerto Morelos, Mexico – (aquarium) | 28° C | Ga0099817_1157661, Ga0099817_1628238 | [30] |
| *Leptolyngbya* sp. SIO1E4 | VI | 28 | Microbial communities of thrombolites from Highborne Cay, Bahamas | 3-5 mm depth | Ga0103786_1000070, Ga0103787_1000070 | [29] |
| | | | *Siderastrea* coral microbial communities from Puerto Morelos, Mexico – (aquarium) | 28° C | Ga0099816_1499140, Ga0099817_1046576, Ga0099817_11128282, Ga0099818_1127966, Ga0099818_1202449, | [30] |
| | | | | 32° C | Ga0099819_1057501, Ga0099819_1081719, Ga0099820_1068013, Ga0099820_1093072, Ga0099821_1042070, Ga0099821_1506664 | [30] |

**Table S10.** Metatranscriptomics datasets from the IMG/MER database. Sequences from three environments can be assigned to three phylogenetic groups (I, IV, VI). All FaRLiP cluster shapes are listed in Figure S3.

| Label | Sampling environment | k-mer size | Assembled metagenome size (Gbps) | Number of FaRLiP contigs (>1 CDS) | Largest FaRLiP –containing contig (bps) | Associated research |
|---|---|---|---|---|---|---|
| **at** | Endolith, rocks, Atacama Desert, Chile | 119 | 633 | 5 | 16,687 | [31] |
| **bah** | Thrombolites, Bahamas | 129 | 59 | 6 | 12,059 | [32] |
| **cl** | Thrombolites, Clifton Lake, Australia | 119 | 738 | 11 | 122,739 | [6] |
| **mexn** | Stromatolites, Alchichica Lake (North), Mexico | 117 | 3,276 | 35 | 68,907 | [5] |
| **mexw** | Stromatolites, Alchichica Lake (West), Mexico | 117 | 2,369 | 26 | 68,907 | -idem- |
| **pac2** | Pacific Garbage Patch, sampling point #2 | 141 | 1,587 | 48 | 40,066 | [7] |
| **pac5** | Pacific Garbage Patch, sampling point #5 | 141 | 2,534 | 34 | 24,636 | -idem- |
| **pac9** | Pacific Garbage Patch, sampling point #9 | 141 | 2,775 | 44 | 71,134 | -idem- |
| **pac11** | Pacific Garbage Patch, sampling point #11 | 141 | 872 | 46 | 91,565 | -idem- |
| **pac14** | Pacific Garbage Patch, sampling point #14 | 141 | 2,604 | 44 | 32,298 | -idem- |
| **pac15** | Pacific Garbage Patch, sampling point #15 | 141 | 1,701 | 34 | 8,338 | -idem- |
| **rivr** | River estuary, Elkhorn Slough, CA, USA | 141 | 814 | 0 | N/A | |
| **sbc** | Shark Bay stromatolites (colloform), Australia | 127 | 3,232 | 9 | 3,032 | [8] |
| **sbp** | Shark Bay stromatolites (pustular), Australia | 127 | 2,283 | 7 | 17,243 | -idem- |
| **sbs** | Shark Bay stromatolites (smooth), Australia | 127 | 2,352 | 6 | 3,991 | -idem- |

**Table S12.** Metagenome co-assembly details. Most of the datasets were assembled with the 'meta-large' preset setting due to their size. A subset (k-mer size ending in 9) used 'meta-sensitive'. The FaRLiP custer covers ~25 kbps, and is sometimes split in two. Many of the assemblies were unable to recover a full cluster, but few included large contigs clearly placing FaRLiP genes within a much larger gene neighborhood (see Clifton Lake data statistics). The 'rivr' dataset contained and *apcE2* fragment, but a full cluster could not be assembled. Overviews of the metagenomes and their associated environments are listed.

| Function | Identifier | Gene | | Protein | |
|---|---|---|---|---|---|
| | | Ambiguity index | Root Point | Ambiguity index | Root Point |
| Phycobilisome | *apcB2* | 0.959 | b | 0.968 | - |
| | *apcD2* | 0.992 | b | 0.990 | c |
| | *apcD3* | **0.844** | a* | 0.938 | a* |
| | *apcD5* | 0.993 | - | 1.000 | - |
| | *apcE2* | **0.902** | a | 0.928 | a |
| Photosystem I | *psaA2* | 0.977 | b* | 0.996 | c* |
| | *psaB2* | 0.997 | c | 0.935 | c |
| | *psaF2* | 0.928 | b | 0.994 | b |
| | *psaI2* | 0.996 | a | 0.983 | a* |
| | *psaJ2* | 0.987 | c | 0.979 | - |
| | *psaL2* | 0.914 | c | 0.971 | - |
| Chl *f* synthase | *chlF* | 0.986 | a | 0.949 | a |
| Photosystem II | *psbA3* | 0.984 | - | 0.996 | - |
| | *psbB2* | 0.971 | b | 0.929 | - |
| | *psbC2* | **0.820** | a | **0.821** | a |
| | *psbD3* | 1.000 | a | 0.926 | a |
| Phytochrome signaling cascade | *rfpA* | 0.994 | a | 0.969 | a |
| | *rfpB* | 0.977 | a | 0.983 | a |
| | *rfpC* | 0.997 | a | 0.991 | a |

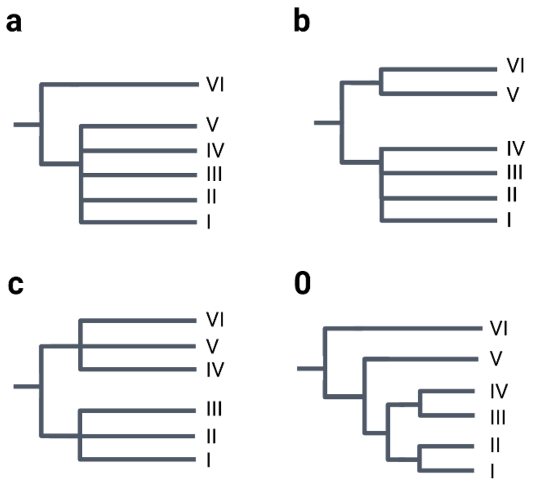


**Table S13.** Root point identification with Minimal Ancestral Deviation (MAD). FaRLiP gene and protein trees exhibited a range of root points. The most common ones were labeled as follows: ‘a’ (group VI as outgroup, 18 entries), ‘b’ (groups V and VI as outgroups, 6 entries), and ‘c’ (split between group IV-VI and I-III, 6 entries). These topologies are shown on the right. Rarer versions are marked with dashes. Root point topology ‘a’ is not only the most common version, but it is the version associated with the most reliable root point splits (lowest Ambiguity Indices, marked in bold). This suggests it is the true root point, with others being artifacts.

| Current work | Number | GTDB Taxonomy | NCBI Taxonomy and 16S rRNA-based literature | Recent literature[33] |
|---|---|---|---|---|
| **Gloeobacterales** | (root) | Gloeobacterales | Gloeobacterales | Gloeobacterales |
| **Nodosilineales** | VI | Phormidesmiales | Prochlorotrichaceae, Synechococcales | Nodosilineales |
| **Oculatellales** | V | Elainellales | Oculatellaceae, Synechococcales | Oculatellales |
| **Leptolyngbyales** | V | Leptolyngbyales | Leptolyngbyaceae, Synechococcales | Leptolyngbyales |
| **Chroococcales** | IV | Cyanobacteriales (Microcystaceae/Hydrococcus and Xenococcaceae) | Pleurocapsales | Chroococcales (Microcystaceae and Pleurocapsaceae) |
| **Coleofasciculales** | III | Cyanobacteriales (genome not available) | Coleofasciculaceae, Oscillatoriales | Coleofasciculales |
| **Chroococcidiopsidales** | II | Cyanobacteriales (Chroococcidiopsidaceae) | Chroococcidiopsidales | Chroococcidiopsidales |
| **Nostocales** | I | Cyanobacteriales (Nostocaceae) | Nostocales | Nostocales |

**Table S14**. Taxonomic definitions employed in this paper. The Genome Taxonomy Database (GTDB) uses whole genome/metagenome information to classify strains. The NCBI Taxonomy Database uses 16S rRNA [34]. We used GTDB-Tk to classify the genomes and metagenomes in this study, but updated the naming where required[33].

### Supplemental references


1. Billi, D., Napoli, A., Mosca, C., Fagliarone, C., Carolis, R. De, Balbi, A., Scanu, M., Selinger, V.M., Antonaru, L.A., and Nürnberg, D.J. (2022). Identification of far-red light acclimation in an endolithic Chroococcidiopsis strain and associated genomic features : Implications for oxygenic photosynthesis on exoplanets. Front. Microbiol. *13*, 1–12. 10.3389/fmicb.2022.933404.
2. CCAP (2023). JM (Jaworski's Medium). https://www.ccap.ac.uk/wp-content/uploads/MR_JM.pdf.
3. CCAP (2021). ASW:BG medium for marine cyanobacteria. https://www.ccap.ac.uk/wp-content/uploads/MR_ASW_BG.pdf.
4. Rippka, R., Deruelles, J., Waterbury, J.B., Herdman, M., and Stanier, R.Y. (1979). Generic assignments, strain histories and properties of pure cultures of cyanobacteria. J. Gen. Microbiol. *111*, 1–61. 10.1099/00221287-111-1-1.
5. Saghaï, A., Zivanovic, Y., Moreira, D., Benzerara, K., Bertolino, P., Ragon, M., Tavera, R., López-Archilla, A.I., and López-García, P. (2016). Comparative metagenomics unveils functions and genome features of microbialite-associated communities along a depth gradient. Environ. Microbiol. *18*, 4990–5004. 10.1111/1462-2920.13456.
6. Warden, J.G., Casaburi, G., Omelon, C.R., Bennett, P.C., Breecker, D.O., and Foster, J.S. (2016). Characterization of microbial mat microbiomes in the modern thrombolite ecosystem of lake Clifton, Western Australia using shotgun metagenomics. Front. Microbiol. *7*, 1–14. 10.3389/fmicb.2016.01064.
7. Bryant, J.A., Clemente, T.M., Viviani, D.A., Fong, A.A., Thomas, K.A., Kemp, P., Karl, D.M., White, A.E., and DeLong, E.F. (2016). Diversity and Activity of Communities Inhabiting Plastic Debris in the North Pacific Gyre. mSystems *1*, 1–19. 10.1128/msystems.00024-16.
8. Babilonia, J., Conesa, A., Casaburi, G., Pereira, C., Louyakis, A.S., Reid, R.P., and Foster, J.S. (2018). Comparative metagenomics provides insight into the ecosystem functioning of the Shark Bay Stromatolites, Western Australia. Front. Microbiol. *9*, 1–16. 10.3389/fmicb.2018.01359.
9. Murray, B., Ertekin, E., Dailey, M., Soulier, N.T., Shen, G., Bryant, D.A., Perez-Fernandez, C., and Diruggiero, J. (2022). Adaptation of Cyanobacteria to the endolithic light spectrum in hyper-arid deserts. Microorganisms *10*, 1–12. 10.3390/microorganisms10061198.
10. Finke, N., Simister, R.L., O'Neil, A.H., Nomosatryo, S., Henny, C., MacLean, L.C., Canfield, D.E., Konhauser, K., Lalonde, S. V., Fowle, D.A., et al. (2019). Mesophilic microorganisms build terrestrial mats analogous to Precambrian microbial jungles. Nat. Commun. *10*, 1–12. 10.1038/s41467-019-11541-x.
11. Brawley, S.H., Blouin, N.A., Ficko-Blean, E., Wheeler, G.L., Lohr, M., Goodson, H. V., Jenkins, J.W., Blaby-Haas, C.E., Helliwell, K.E., Chan, C.X., et al. (2017). Insights into the red algae and eukaryotic evolution from the genome of Porphyra umbilicalis (Bangiophyceae, Rhodophyta). Proc. Natl. Acad. Sci. U. S. A. *114*, E6361–E6370. 10.1073/pnas.1703088114.
12. Antonaru, L.A., Cardona, T., Larkum, A.W.D., and Nürnberg, D.J. (2020). Global distribution of a chlorophyll f cyanobacterial marker. ISME J. *14*, 2275–2287. 10.1038/s41396-020-0670-y.
13. Miao, D., Ding, W.L., Zhao, B.Q., Lu, L., Xu, Q.Z., Scheer, H., and Zhao, K.H. (2016). Adapting photosynthesis to the near-infrared: Non-covalent binding of phycocyanobilin provides an extreme spectral red-shift to phycobilisome core-membrane linker from Synechococcus sp. PCC7335. Biochim. Biophys. Acta - Bioenerg. *1857*, 688–694. 10.1016/j.bbabio.2016.03.033.
14. Trinugroho, J.P., Bečková, M., Shao, S., Yu, J., Zhao, Z., Murray, J.W., Sobotka, R., Komenda, J., and Nixon, P.J. (2020). Chlorophyll f synthesis by a super-rogue photosystem II complex. Nat. Plants *6*, 238–244. 10.1038/s41477-020-0616-4.
15. Gisriel, C.J., Shen, G., Ho, M.-Y.Y., Kurashov, V., Flesher, D.A., Wang, J., Armstrong, W.H., Golbeck, J.H., Gunner, M.R., Vinyard, D.J., et al. (2022). Structure of a monomeric photosystem II core complex from a cyanobacterium acclimated to far-red light reveals the

functions of chlorophylls d and f. J. Biol. Chem. *298*, 1–15. 10.1016/j.jbc.2021.101424.
16. Nürnberg, D.J., Morton, J., Santabarbara, S., Telfer, A., Joliot, P., Antonaru, L.A., Ruban, A. V., Cardona, T., Krausz, E., Boussac, A., et al. (2018). Photochemistry beyond the red limit in chlorophyll f–containing photosystems. Science *360*, 1210–1213. 10.1126/science.aar8313.
17. Gisriel, C.J., Cardona, T., Bryant, D.A., and Brudvig, G.W. (2022). Molecular Evolution of Far-Red Light-Acclimated Photosystem II. Microorganisms *10*, 1–18. 10.3390/microorganisms10071270.
18. Gisriel, C.J., Flesher, D.A., Shen, G., Wang, J., Ho, M.Y., Brudvig, G.W., and Bryant, D.A. (2022). Structure of a photosystem I-ferredoxin complex from a marine cyanobacterium provides insights into far-red light photoacclimation. J. Biol. Chem. *298*, 101408. 10.1016/j.jbc.2021.101408.
19. Ho, M.Y., and Bryant, D.A. (2019). Global transcriptional profiling of the cyanobacterium Chlorogloeopsis fritschii PCC 9212 in far-red light: Insights into the regulation of chlorophyll d synthesis. Front. Microbiol. *10*, 1–16. 10.3389/fmicb.2019.00465.
20. Yan, J., and Kurgan, L. (2017). DRNApred, fast sequence-based method that accurately predicts and discriminates DNA-and RNA-binding residues. Nucleic Acids Res. *45*, 1–16. 10.1093/nar/gkx059.
21. Marchler-Bauer, A., Bo, Y., Han, L., He, J., Lanczycki, C.J., Lu, S., Chitsaz, F., Derbyshire, M.K., Geer, R.C., Gonzales, N.R., et al. (2017). CDD/SPARCLE: Functional classification of proteins via subfamily domain architectures. Nucleic Acids Res. *45*, D200–D203. 10.1093/nar/gkw1129.
22. Komenda, J., Tichý, M., and Eichacker, L.A. (2005). The PsbH protein is associated with the inner antenna CP47 and facilitates D1 processing and incorporation into PSII in the cyanobacterium Synechocystis PCC 6803. Plant Cell Physiol. *46*, 1477–1483. 10.1093/pcp/pci159.
23. Popelkova, H., and Yocum, C.F. (2011). PsbO, the manganese-stabilizing protein: Analysis of the structure-function relations that provide insights into its role in photosystem II. J. Photochem. Photobiol. B Biol. *104*, 179–190. 10.1016/j.jphotobiol.2011.01.015.
24. Barber, J. (2016). "Photosystem II: the water splitting enzyme of photosynthesis and the origin of oxygen in our atmosphere." Q. Rev. Biophys. *49*, 1–21. 10.1017/s0033583516000093.
25. Müh, F., Renger, T., and Zouni, A. (2008). Crystal structure of cyanobacterial photosystem II at 3.0 Å resolution: A closer look at the antenna system and the small membrane-intrinsic subunits. Plant Physiol. Biochem. *46*, 238–264. 10.1016/j.plaphy.2008.01.003.
26. Chen, M., Hernandez-Prieto, M.A., Loughlin, P.C., Li, Y., and Willows, R.D. (2019). Genome and proteome of the chlorophyll f-producing cyanobacterium Halomicronema hongdechloris: Adaptative proteomic shifts under different light conditions. BMC Genomics *20*, 1–15. 10.1186/s12864-019-5587-3.
27. MacGregor-Chatwin, C., Nürnberg, D.J., Jackson, P.J., Vasilev, C., Hitchcock, A., Ho, M.Y., Shen, G., Gisriel, C.J., Wood, W.H.J., Mahbub, M., et al. (2022). Changes in supramolecular organization of cyanobacterial thylakoid membrane complexes in response to far-red light photoacclimation. Sci. Adv. *8*, 1–16. 10.1126/sciadv.abj4437.
28. Ho, M.Y., Gan, F., Shen, G., Zhao, C., and Bryant, D.A. (2017). Far-red light photoacclimation (FaRLiP) in Synechococcus sp. PCC 7335: I. Regulation of FaRLiP gene expression. Photosynth. Res. *131*, 173–186. 10.1007/s11120-016-0309-z.
29. Mobberley, J.M., Khodadad, C.L.M., Visscher, P.T., Reid, R.P., Hagan, P., and Foster, J.S. (2015). Inner workings of thrombolites: Spatial gradients of metabolic activity as revealed by metatranscriptome profiling. Sci. Rep. *5*, 1–15. 10.1038/srep12601.
30. Avila-Magaña, V., Kamel, B., DeSalvo, M., Gómez-Campo, K., Enríquez, S., Kitano, H., Rohlfs, R. V., Iglesias-Prieto, R., and Medina, M. (2021). Elucidating gene expression adaptation of phylogenetically divergent coral holobionts under heat stress. Nat. Commun. *12*, 1–16. 10.1038/s41467-021-25950-4.
31. Crits-Christoph, A., Robinson, C.K., Ma, B., Ravel, J., Wierzchos, J., Ascaso, C., Artieda, O., Souza-Egipsy, V., Casero, M.C., and DiRuggiero, J. (2016). Phylogenetic and functional

substrate specificity for endolithic microbial communities in hyper-arid environments. Front. Microbiol. *7*, 1–15. 10.3389/fmicb.2016.00301.

32. Louyakis, A.S., Mobberley, J.M., Vitek, B.E., Visscher, P.T., Hagan, P.D., Reid, R.P., Kozdon, R., Orland, I.J., Valley, J.W., Planavsky, N.J., et al. (2017). A Study of the Microbial Spatial Heterogeneity of Bahamian Thrombolites Using Molecular, Biochemical, and Stable Isotope Analyses. Astrobiology *17*, 413–430. 10.1089/ast.2016.1563.
33. Strunecký, O., Ivanova, A.P., and Mareš, J. (2022). An updated classification of cyanobacterial orders and families based on phylogenomic and polyphasic analysis. J. Phycol. *51*, 12–51. 10.1111/jpy.13304.
34. Dextro, R.B., Delbaje, E., Cotta, S.R., Zehr, J.P., and Fiore, M.F. (2021). Trends in Free-access Genomic Data Accelerate Advances in Cyanobacteria Taxonomy. J. Phycol. *57*, 1392–1402. 10.1111/jpy.13200.
